# LRPPRC gene replacement therapy preserves cardiac function and prevents premature death in a mouse model of mitochondrial disease

**DOI:** 10.64898/2026.11.12.762002

**Authors:** Sofia Zanin, Franck Lager, Marcelo Simon-Sola, Clovis Gaborit, Gilles Renault, Federico Mingozzi, Agnes Rotig, Marina Cavazzana, Metodi D. Metodiev, Benedetta Ruzzenente

## Abstract

Pathogenic variants in LRPPRC disrupt mitochondrial gene expression, causing severe and incurable mitochondrial disease with neurological involvement accompanied by hepatopathy or cardiomyopathy. The development for gene therapy for LRPPRC-linked disease is hindered by the large LRPPRC coding sequence exceeding AAV cargo capacity when combined with classical regulatory elements. Here, we engineered compact single rAAV9 vectors expressing human or mouse LRPPRC and administered them intravenously to heart and skeletal muscle-specific *Lrpprc* knockout mice. This approach restored cardiac LRPPRC expression, preserved oxidative phosphorylation (OXPHOS) function, prevented cardiomyopathy and markedly extended survival. While skeletal muscle rescue was partial, the therapy effectively prevented fatal cardiac dysfunction. These findings provide the first in vivo proof-of-concept that AAV-mediated LRPPRC gene replacement can preserve OXPHOS function and prevent lethal cardiomyopathy, supporting its potential as a therapeutic strategy for mitochondrial disease linked to LRPPRC deficiency.

## Introduction

Mitochondrial diseases (MD) are among the most frequent metabolic disorders caused by defects in oxidative phosphorylation (OXPHOS), the main process responsible for ATP production by the so-called OXPHOS system. The latter consists of five multisubunit complexes (CI-CV), four of which (CI, CIII-CV) contain subunits encoded by nuclear and mitochondrial DNA (mtDNA). Because these complexes have a dual genetic origin, mutations in either genome can lead to MD. To date, pathogenic variants have been identified in all 37 mitochondrial genes and in nearly 400 nuclear genes [1]. Collectively, MDs have an estimated frequency of ∼1:5000 [2], although when each individual gene is considered, they are even rarer, sometimes with only a few clinical cases per gene worldwide. Besides their genetic heterogeneity, MD are also clinically heterogeneous, where mutations in the same gene can cause different clinical presentations. Because mitochondria are ubiquitous, most tissues and organs can be implicated in MD, yet these pathologies can have unexplained tissue specificity. Many genes involved in MD are key regulators of mitochondrial genome expression [3]. Their function spans all steps of this process, from mtDNA maintenance and replication to the translation and assembly of the 13 mtDNA encoded OXPHOS subunits. Mutations in these genes typically cause multiple OXPHOS deficiencies with two or more dual-origin complexes affected, while complex II, which is exclusively nucleus-encoded, is spared.

The *LRPPRC* gene encodes the Leucine Rich Pentatricopeptide Repeat-Containing Protein, LRPPRC, an important mitochondrial mRNA stability and translation factor [4]. Pathogenic *LRPPRC* variants have been linked to neurological disease, including Leigh syndrome and leukoencephalopathy, accompanied by hepatopathy or cardiomyopathy [5–12]. Disease onset is usually between birth and five months of age, with most severely affected children succumbing to a fatal metabolic crisis before the age of two. To date, numerous patients with pathogenic *LRPPRC* variants have been reported worldwide. Most variants are extremely rare, except the A354V variant which, due to a founder effect in the Quebec region of Canada, has a carrier frequency of ∼1 in 23 and impacts ∼1 in 2000 live births in this region [6]. Because of this, the syndrome caused by the A354V variant is called Leigh Syndrome French Canadian Variant (LSFC). Biochemically, the tested pathogenic variants all lead to decreased LRPPRC levels in patient biopsies and fibroblasts [12–15]. Moreover, heart, skeletal muscle, and liver biopsies show OXPHOS complex IV defects, whereas patient fibroblasts and myoblasts present in addition variable complex I and V deficiencies [12–14]. These findings are consistent with observations from two tissue-specific knockout mouse models: (1) a heart and skeletal muscle *Lrpprc* knockout model exhibiting reduced lifespan and profound deficiency in complex IV and complex V [4], and (2) a liver specific knockout model showing no change in survival, despite growth delay, hepatic tissue damage and reduced complex IV and V activities [16]. Mechanistically, loss of LRPPRC leads to mitochondrial mRNAs instability accompanied by shortening of the poly(A) tail and extensive remodeling of mRNA secondary structure [4, 17, 18]. This results in dysregulated mitochondrial translation which impairs OXPHOS biogenesis.

Like all mitochondrial diseases, *LRPPRC*-linked disease is incurable, while available treatments are only symptomatic and unable to prevent the progression of the disease. LRPPRC gene replacement therapy (GRT) holds significant promise for curing *LRPPRC*-linked disease by introducing a wild-type copy of the mutated gene via a viral or non-viral delivery strategy. For viral delivery, adeno-associated virus (AAV) vectors are commonly used because they are non-integrating, have low immunogenicity, and do not cause disease in humans. Several gene replacement therapies targeting LHON caused by pathogenic variants in the mtDNA-encoded ND1 and ND4 genes are currently in clinical trials. In addition, there are several ongoing clinical trials targeting the *FRATAXIN* gene implicated in Friedreich’s ataxia. Despite the large number of nuclear and mtDNA-encoded genes linked to MD, proof-of-concept studies for GRT have been reported for only ∼17 genes. This limited progress stems from several challenges, including the small number of patients per gene, the multisystemic nature of mitochondrial disease, the need to target multiple tissues, and the limited availability of suitable animal models for proof-of-concept studies. In many cases, whole-body knockouts of mitochondrial proteins are embryonic lethal, while knock-in mice do not always develop a physiological phenotype with clear-cut therapeutic endpoints. This further reduces the number of available models or mandates the use of tissue-specific knockout mice, limiting proof-of-concept studies to one or a few organs at a time. MtDNA-encoded genes present an additional complication, as their protein products, synthesized in mitochondria, are typically very hydrophobic, making cytosol expression (allotopic expression) challenging. Finally, the limited cargo capacity of AAV (less than 4.7-5 kb including all regulatory elements) has hindered the development of GRT for large genes such as *LRPPRC*. Although different dual-vector strategies offer potential solutions [19], they come at the cost of reduced transduction efficiency and low or mosaic transgene expression because two vectors, each carrying half of the coding sequence, must co-transduce the same cell. Consequently, GRT development in the MD field has focused primarily on relatively small genes, with the mitochondrial dynamin-like GTPase OPA1 (∼2994 bp) being the largest transgene tested but still within the AAV cargo capacity [20]. Proof-of-concept studies have been reported for the OXPHOS subunits NDUFS3, NDUFS4, NDUFS6, COX10 [21–25], the OXPHOS assembly factors (SURF1 and BCS1L) [26, 27], mitochondrial transporters (ANT, SLC25A46) [28, 29], mitochondrial metabolic enzymes involved in various pathways (ETHE1, MPV17, TK2, TYMP, TAFFAZIN, GTPBP3, DGUOK) [30–38], FRATAXIN involved in iron metabolism [39, 40] and GFM1 involved in mitochondrial translation [41].

Here, we present the first in vivo proof-of-concept for LRPPRC gene replacement therapy. Because the LRPPRC cDNA is large (4185bp), we engineered rAAV9 vectors incorporating only minimal and essential regulatory sequences. We showed that a single-dose injection of optimized rAAV9 viral particles expressing hLRPPRC or mLRPPRC in heart- and skeletal-muscle-specific knockout mice provided a clear therapeutic benefit by preserving mitochondrial function in the heart, preventing cardiomyopathy and extending lifespan. Moreover, this optimized single-vector approach demonstrated superior therapeutic efficacy compared with the conventional trans-splicing dual-vector strategy with standard regulatory elements.

## Results

### Mouse model and therapeutic endpoints

Currently, there are no mouse models that simultaneously recapitulate the main neurological, hepatological and cardiac features of *LRPPRC* disease, and whole-body *Lrpprc* knockout is embryonic lethal [4]. Therefore, to develop a proof-of-concept for *LRPPRC* GRT, *in vivo*, we used a heart- and-skeletal muscle-specific *Lrpprc* knockout (henceforth KO) mouse model that we previously developed and characterized [4]. In this model, Cre recombinase was expressed under the control of the muscle creatine kinase promoter (*ckmm*), leading to *Lrpprc* ablation in cardiomyocytes and skeletal muscle fibers. Using a transgenic mouse line carrying a Cre-inducible tdTomato reporter, we confirmed that Cre expression, and thus *Lrpprc* inactivation, occurs simultaneously in cardiomyocytes and in all skeletal muscle fibers (Figure S1A). Consistent with our previous reports, KO mice developed progressive cardiomyopathy starting after 8 weeks of age. In addition to the previously reported increase in heart-to-body weight ratio, echocardiography revealed a significant decrease in ejection fraction and an increase in left ventricular diastolic volume in near-terminal mice, establishing clear pathophysiological markers for evaluating the therapeutic benefit of *LRPPRC* GRT (Figure S1B). KO mice died from heart failure between 7 and 13 weeks (Figure S1C). Although skeletal muscle dysfunction has not been previously characterized in this model, lethality is confidently attributed to cardiomyopathy rather than skeletal muscle involvement.

### Low therapeutic efficiency of a trans-splicing dual vector for *LRPPRC* gene therapy

To overcome the challenge posed by the large size of *LRPPRC* cDNA, initially, we adopted a trans-splicing dual-vector strategy, which has been successfully used for other large transgenes [42, 43]. We designed two vectors: the first containing a CMV promoter followed by a compact chimeric intron (from the human β-globin and the immunoglobulin heavy chain genes) and the first 20 exons of *LRPPRC* fused to a splice donor sequence whereas the second plasmid contained a splice acceptor sequence linked to the remaining 18 exons fused to a FLAG tag, followed by the Woodchuck Hepatitis Virus Posttranscriptional Regulatory Element (WPRE) sequence and the bovine growth hormone polyadenylation (polyA) signal (Figure S1D). We previously demonstrated that a C-terminal FLAG tag does not impair the Lrpprc function *in vivo* by rescuing the lethality of the whole-body *Lrpprc* knockout [4]. An important limitation of the dual vector strategy is that formation of full-length hLRPPRC requires co-transduction of both vectors in the same cell, correct vector orientation, and accurate splicing between the donor and acceptor sites, factors that can limit overall transgene expression efficiency. Next, we generated rAAV9 particles from either plasmid. We chose this capsid because of its tissue-wide tropism and the multisystemic nature of *LRPPRC*-linked disease. To assess hLRPPRC expression and therapeutic efficacy, we injected 3-to 4-week-old KO mice intravenously into the caudal vein with either the dual-vector system (1:1 vector mix) or a GFP-expressing control vector at a total dose of 2 × 10¹³ vg/kg (Figure S1E). Some treated KO mice were sacrificed at 11 weeks (the estimated terminal age for untreated KO mice), while others were retained to assess the effect of this treatment on lifespan. Heart-to-body weight ratio of treated 11-week-old KO mice was lower than that of GFP-injected KO mice, but significantly higher than that of controls, suggesting only a modest therapeutic effect of the dual-vector strategy (data not shown). To assess the expression of full-length hLRPPRC, we carried out immunoblot analysis of heart mitochondrial extracts using anti-LRPPRC antibodies raised against the C-terminal 380 amino acids of hLRPPRC. However, as this region shares 68% sequence identity with the mouse LRPPRC, the antibodies also recognized the mouse protein. This analysis confirmed a modest expression of full-length hLRPPRC (Figure S1F). Immunofluorescent analysis of heart sections showed a limited number of cardiomyocytes expressing hLRPPRC (Figure S1G), consistent with COX histochemistry and Western blot data revealing only marginal preservation of complex IV subunits and activity (Figure S1F and H). Additionally, the lifespan of treated mice was only marginally extended. These results demonstrate that the overall therapeutic benefit of the trans-splicing dual-vector approach for LRPPRC GRT was low. The sporadic expression of LRPPRC throughout the myocardium illustrates the above-mentioned drawbacks of using this approach for LRPPRC GRT.

### Development of a single-vector approach for the AAV-mediated expression of hLRPPRC

Given the low efficiency of the trans-splicing dual-vector strategy, we developed a single-vector approach utilizing only essential minimized regulatory elements to remain within or close to the AAV cargo capacity (Figure 1A). These new vectors incorporated the same ITRs sequences and the compact chimeric intron used in the trans-splicing dual vector strategy. However, they lacked the WPRE sequence, and the bGH polyA signal was replaced with the smaller SV40 polyA signal. We selected two compact, ubiquitously active promoters: miniCMV and the short form of EF1α (EFS) [44, 45]. As before, hLRPPRC carried a C-terminal FLAG tag. This strategy yielded two plasmids: TG5, in which the miniCMV promoter drove hLRPPRC:FLAG expression, and TG7, which carried the EFS promoter instead. The total cassette sizes, including all regulatory elements, were at the limit of AAV packaging capacity (5,080 bp and 5,160 bp, for TG5 and TG7, respectively) (Figure 1A). To generate control plasmids, we replaced hLRPPRC:FLAG with GFP, yielding TG5-GFP and TG7-GFP (Figure 1A). To confirm the ability of these plasmids to express the transgenes, we transiently transfected them in HeLa cells followed by immunofluorescence using FLAG-specific antisera. This analysis showed that both plasmids expressed hLRPPRC, which was correctly targeted to mitochondria, as evidenced by colocalization with the mitochondrial marker TOM20, whereas GFP showed diffuse cytoplasmic or weak nuclear staining because it lacked a mitochondrial localization signal. (Figure 1B).

**Figure 1.**
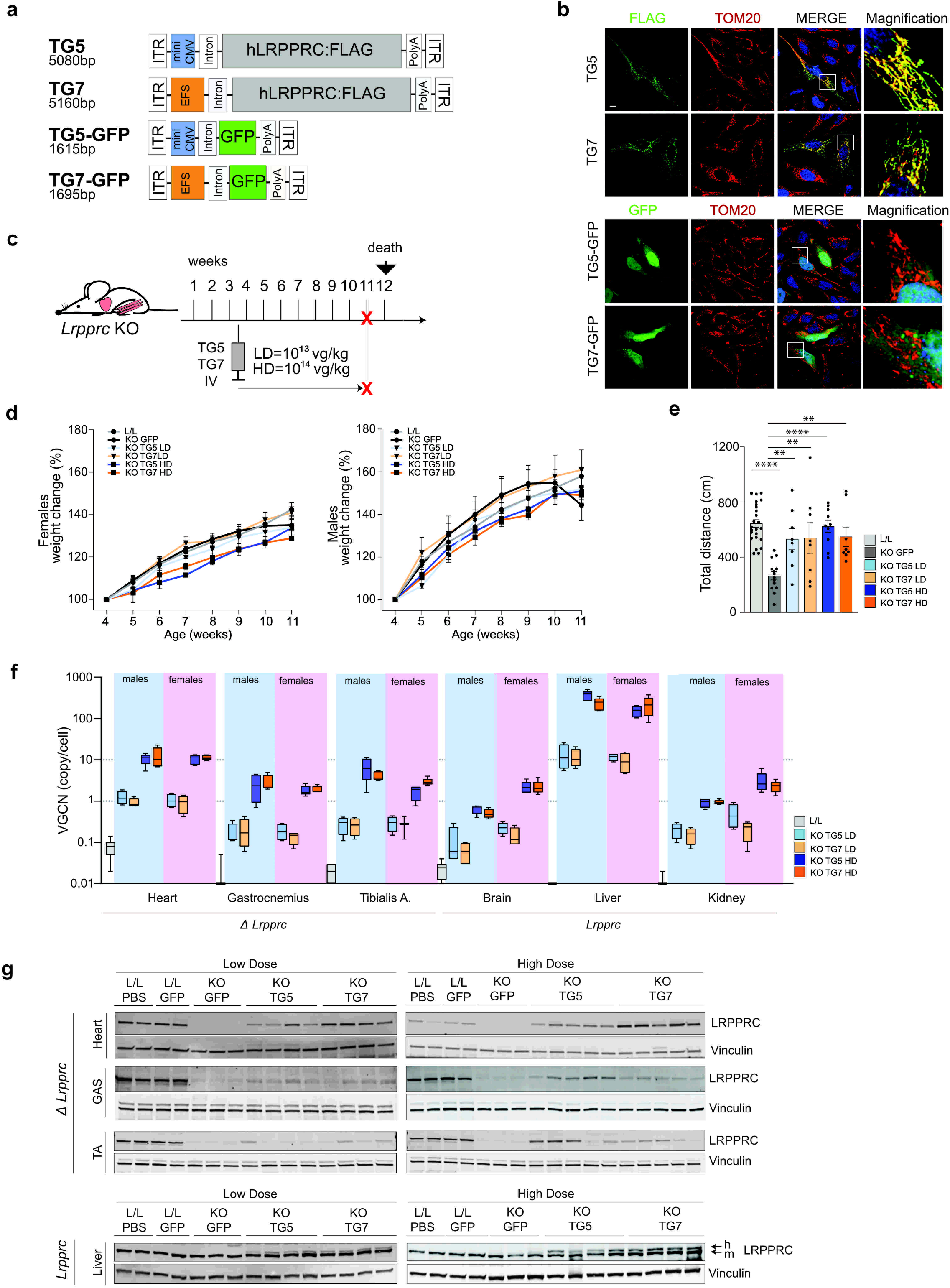
Design of optimized single-rAAV vector constructs for hLRPPRC expression in vitro and in vivo. (A) Schematic representation of the optimized single-vector rAAV9 constructs (TG5 and TG7) expressing hLRPPRC:FLAG and the corresponding GFP control vectors. Sizes between the ITRs are indicated. (B) Representative confocal images of HeLa cells transiently transfected with TG5 or TG7 plasmids, showing hLRPPRC:FLAG expression and its mitochondrial localization. hLRPPRC was detected by FLAG antibody whereas TOM20 was used as a mitochondrial marker. Scale bar:10 µm. (C) Experimental design. 3–4-week-old *Lrpprc* KO mice were intravenously injected with low-dose (LD, 10¹³ vg/kg) or high-dose (HD, 10¹⁴ vg/kg) rAAV9 TG5, TG7, TG5GFP or TG7-GFP. Control mice were instead injected with PBS, TG5-GFP or TG7-GFP at the corresponding LD and HD. Red crosses indicate euthanasia time points. (D) Relative body weight change with age in the different experimental groups. Left, female mice; right, male mice. Data for PBS- or LD and HD GFP-injected L/L mice were pooled. For female mice, L/L, n=12; KO, pooled LD and HD GFP-injected, n=6; KO TG5 LD, n=4; KO TG7 LD, n=4; KO TG5 HD, n=5; KO TG7 HD, n=5. For male mice: L/L, n=12; KO, pooled LD and HD GFP-injected, n=8; KO TG5 LD, n=4; KO TG7 LD, n=4; KO TG5 HD, n=5; KO TG7 HD, n=5. (E) Open-field test of locomotor performance in the different treatment groups. Data for male and female mice were pooled. L/L, n=24; KO, n=14; KO TG5 LD, n=8; KO TG7 LD, n=8; KO TG5 HD, n=10; KO TG7 HD, n=8. One-way ANOVA test with Dunnett’s correction: **, p<0.01; ****, p<0.0001. (F) Vector genome copy number (VGCN) per cell in the indicated tissues with (Δ*Lrpprc*) or without (*Lrpprc*) ablation of endogenous *Lrpprc*. L/L, n=6-8; KO TG5 LD, males and females each n=4; KO TG7 LD, males n=4, females n=3-4; KO TG5 HD, males n=5, females n=4-5; KO TG7 HD, males and females each n=5. (G) Western blot analysis of hLRPPRC protein levels in total tissue protein extracts from the heart, gastrocnemius, tibialis anterior, and liver of male mice from the indicated experimental groups. Vinculin was used as a loading control. Corresponding analyses of female mice are shown in Figure S2A.

Next, we generated rAAV9 viral particles using all four plasmids and assessed the therapeutic potential of TG5 and TG7 in *Lrpprc* KO mice. Presymptomatic KO mice, at 3-4 weeks of age, were injected into the caudal vein with the different vectors at two doses: low dose (LD; 10^13^vg/kg) and high dose (HD; 10^14^vg/kg) (Figure 1C). *Lrpprc^Lox/Lox^* (L/L) mice, used as controls, were injected with TG5-GFP, TG7-GFP or PBS. The change in body weight of all injected mice was recorded weekly until they were sacrificed at 11 weeks of age, the experimental endpoint for untreated KO mice. For the duration of the experiment, all mice continued to gain weight, and only GFP-injected male KO mice exhibited a sharp decrease in body weight at 11 weeks of age, as expected for untreated KO mice (Figure 1D). Although GFP-injected KO females did not display significant weight loss at the terminal time point, they had decreased locomotor activity, like their male counterparts, suggesting late-stage disease (Figure 1E). In contrast, the locomotor activity of TG5- and TG7-injected KO mice was significantly higher compared to GFP-injected KO mice, suggesting preserved physical performance (Figure 1E). Interestingly, locomotor activity in LD-injected KO mice showed greater variability when compared to the HD-injected group.

Next, we analyzed the biodistribution of TG5 and TG7 by measuring viral genome copy number per cell (VGCN) in different tissues (Figure 1F). Using hLRPPRC-specific primers, we detected both TG5 and TG7 in all examined tissues, consistent with the known broad tissue tropism of the AAV9 serotype. Across tissues, HD-injected mice had approximately 10 times more vector genome copies than LD-injected mice, consistent with a dose-dependent effect on viral biodistribution. In line with published literature, we detected the highest VGCN in the liver, followed by the heart, skeletal muscle (gastrocnemius and tibialis anterior), kidney, and, finally, the brain, which showed the lowest VGCN (Figure 1F). Next, we investigated the transgene expression in different tissues using SDS-PAGE and immunoblotting (Figure 1G). As observed for VGCN, hLRPPRC protein increased with the higher injection dose. Interestingly, the highest protein levels were detected in the heart rather than in the liver, despite the higher VGCN counts in the latter. In liver, which retains endogenous murine LRPPRC expression, the transgene product was distinguishable as a more slowly migrating band. Similar results were obtained in both sexes (Figure 1G and Figure S2A).

### rAAV9-hLRPPRC expression preserves cardiac OXPHOS biogenesis and prevents cardiomyopathy

We proceeded to analyze the transgene expression in cardiomyocytes from TG5- and TG7-injected mice by immunofluorescence. This analysis revealed robust, dose-dependent cardiomyocyte transduction, with HD yielding LRPPRC-positive staining in almost all cardiomyocytes within the examined cross-sections (Figure 2A and Figure S2B). Cardiomyopathy is characterized by increased cardiomyocyte cross-section area (CSA), which is linked to tissue damage. Cardiomyocytes CSA measurements using transverse heart sections showed that, while GFP-injected KO mice developed higher CSA than control mice, CSA in TG5 or TG7-treated KO mice at both doses was similar to that in controls (Figure 2B and Figure S2C). These results were consistent with the heart-to-body weight ratio analyses (Figure 2C) and demonstrated that injection with TG5 and TG7 prevents the development of cardiomyopathy at least up to 11 weeks of age.

**Figure 2.**
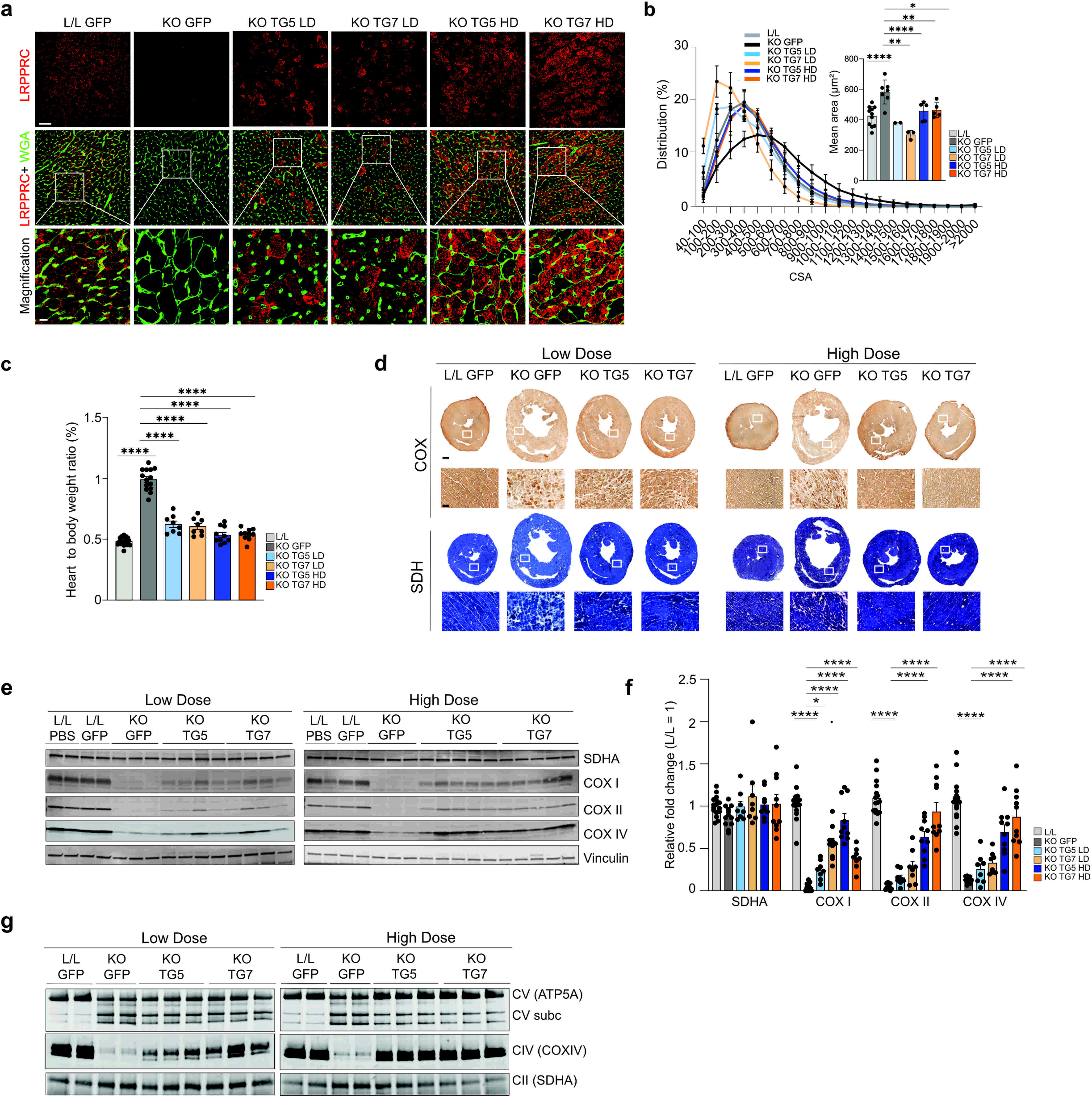
rAAV9-mediated hLRPPRC expression preserves cardiac OXPHOS abundance and prevents cardiomyopathy in KO mice. (A) Representative confocal images of transverse heart sections from 11-week-old male mice demonstrating hLRPPRC expression following administration of the indicated vectors. WGA was used to visualize cardiomyocyte boundaries. Scale bars: main image 50 µm, magnification 10 µm. Corresponding analyses of female mice are shown in Figure S2B. (B) Distribution of cardiomyocyte cross-sectional area (CSA) in heart sections from male mice in the indicated experimental groups. The bar plot shows the mean muscle fiber area. L/L, n=11; KO GFP, n=7; KO TG5 LD, n=2-3; KO TG7 LD, n=3; KO TG5 HD, n=5, KO TG7 HD, n=5. One-way ANOVA test with Dunnett’s correction: *, p<0.05; **, p<0.01; ****, p<0.0001. Corresponding analyses of female mice are shown in Figure S2C. (C) Heart weight-to-body weight ratio in the different experimental groups. Data for male and female mice were pooled. L/L, n=24; KO GFP, n=14; KO TG5 LD, n=8; KO TG7 LD, n=8; KO TG5 HD, n=10, KO TG7 HD, n=10. One-way ANOVA test with Dunnett’s correction: ****, p<0.0001 (D) Representative images of COX/SDH histochemical staining of transverse heart sections for male mice in the different experimental groups. Scale bar: whole section, 500 µm; magnified region, 50 µm. Corresponding analyses of female mice are shown in Figure S2D. (E) Western blot analysis of the steady-state levels of SDHA, COX I, COX II and COX IV proteins in heart homogenates from males in the indicated experimental groups. Vinculin was used as a loading control. Corresponding analyses of female mice are shown in Figure S2E. (F)Quantification of the immunoblots in 2E and S2E. Measurements were pooled within each experimental group. L/L, n=16; KO GFP, n=12; KO TG5 LD, n=8; KO TG7 LD, n=8; KO TG5 HD, n=10, KO TG7 HD, n=10. One-way ANOVA test with Dunnett’s correction: *, p<0.05; **,p<0.01; ****, p<0.0001. (G) BN-PAGE analysis of OXPHOS complexes II, IV and V in heart mitoplast extracts from male mice in the different experimental groups. Corresponding analyses of female mice are shown in Figure S2F.

We previously showed that cardiomyopathy in *Lrpprc* KO mice is driven by defects in OXPHOS biogenesis, with complex IV and complex V being the most affected [4]. Therefore, we analyzed these two complexes using complementary approaches. Histochemical staining of heart transverse sections for complex IV (COX) and complex II (SDH) activity revealed distinct features between GFP- and TG5- or TG7-injected KO mice. As expected, GFP-injected KO hearts exhibited a characteristic dilated left ventricle and strongly reduced COX activity staining (Figure 2D and Figure S2D), whereas complex II activity was similarly intense in the different experimental groups. In contrast, heart sections from TG5- and TG7-injected KO mice revealed normal heart size, absence of left ventricular dilation or fibrosis, and preserved complex IV activity. This was further supported by immunoblot analyses of heart homogenates, which showed that the complex IV subunits, COXI, COXII and COXIV, accumulated in TG5- and TG7-injected mice in a dose-dependent manner (Figure 2E,F and Figure S2E). Finally, BN-PAGE analyses revealed that the abundance of assembled complex IV was preserved in the hearts of TG5- or TG7-injected KO mice, but not in GFP-injected mice (Figure 2G and Figure S2F). Moreover, the accumulation of CV F1 subcomplexes was more pronounced in GFP-injected KO mice but less so in HD TG5- and TG7-injected KO mice.

### Modest LRPPRC expression and partial biochemical rescue in skeletal muscle of TG5- and TG7-injected KO mice

Since *LRPPRC* is also deleted in skeletal muscle of KO mice, we investigated whether there was OXPHOS deficiency and sign of myopathy and if TG5- and TG7-mediated LRPPRC expression can prevent the development of mitochondrial dysfunction in this tissue at 11 weeks of age. We focused on the tibialis anterior and gastrocnemius muscles, which differ in fiber type composition. The tibialis anterior is primarily composed of fast-twitch glycolytic fibers, whereas the gastrocnemius contains a mix of fast- and slow-twitch (oxidative) fibers. After injection with TG5 and TG7, hLRPPRC protein accumulated in these two muscles in a dose-dependent manner, though at seemingly lower levels than in the heart (Figure 1G). Immunofluorescence analysis of sections from both muscle types corroborated the immunoblot analysis: peripheral LRPPRC staining in muscle fibers from TG5- and TG7-injected mice was increased in a dose-dependent manner, whereas it was decreased in untreated KO mice (Figure 3A and Figure S3A). To assess potential muscle damage, we measured muscle fiber cross-sectional areas, but found no differences between control and KO mice injected with the different vectors (Figure 3B). These findings indicate that at 11 weeks of age, the skeletal muscle of *Lrpprc* KO mice shows no overt signs of myopathy. Next, we performed COX/SDH immunohistochemical analyses of muscle sections, which revealed that despite the absence of myopathy, complex IV activity in both muscle types from KO mice injected with different vectors was qualitatively reduced compared with the control group, regardless of vector or injection dose (Figure 3C). These findings were supported by immunoblot analysis of muscle homogenates, which showed reduced levels of COXI, COXII and COXIV proteins in GFP or TG5- or TG7-injected KO mice compared to control mice (Figure 3D,E and Figure S3B).

**Figure 3.**
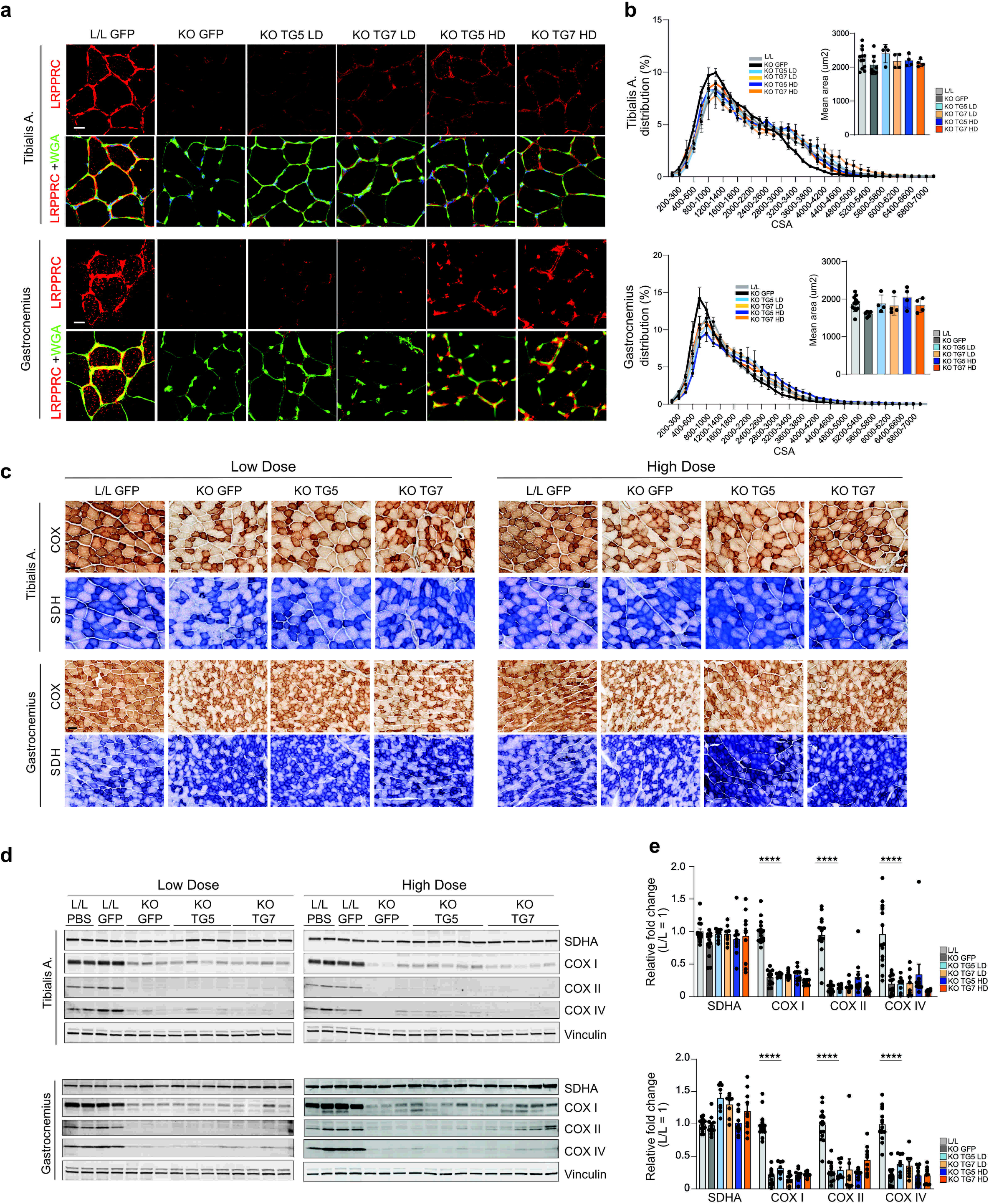
Expression of hLRPPRC does not prevent development of OXPHOS deficiency in skeletal muscle of TG5- and TG7-injected KO mice. (A) Representative confocal images demonstrating hLRPPRC expression in tibialis anterior and gastrocnemius muscles from male mice in the different treatment groups. WGA was used to visualize muscle fiber boundaries. Scale bar: 50 µm. (B) Distribution of muscle fiber CSA in tibialis anterior (top) and gastrocnemius (bottom) muscles. The bar plots show the mean fiber area. L/L, n=12; KO GFP, n=6-8; KO TG5 LD, n=4; KO TG7 LD, n=4; KO TG5 HD, n=4, KO TG7 HD, n=4. (C) Representative images of COX/SDH histochemical staining in transverse skeletal muscle sections from the different treatment groups. Scale bar: 50 µm. (D) Western blot analysis of SDHA, COX I, COX II and COX IV protein abundance in tibialis (top) and gastrocnemius (bottom) homogenates from male mice in the different treatment groups. Corresponding analyses of female mice are shown in Figure S3B. (E) Quantification of the immunoblots in 3D and S3B. Measurements were pooled within each experimental group. L/L, n=15-16; KO GFP, n=12; KO TG5 LD, n=8; KO TG7 LD, n=8; KO TG5 HD, n=10, KO TG7 HD, n=10. One-way ANOVA test with Dunnett’s correction: ****, p<0.0001.

Collectively, these results indicate that TG5- and TG7-mediated hLRPPRC expression prevents the development of OXPHOS dysfunction and cardiomyopathy in *Lrpprc* KO mice. In contrast, transduction in skeletal muscle appears less efficient than in cardiac tissue and insufficient to fully prevent OXPHOS defects. Nonetheless, at this stage, the biochemical defect in skeletal muscle does not yet translate into detectable muscle pathology.

### Stable long-term hLRPPRC expression prolonged the survival of *Lrpprc* KO mice by preserving OXPHOS activity and cardiac function

Our short-term study demonstrated the therapeutic potential of both TG5 and TG7 vectors. However, as all experimental groups were sacrificed at 11 weeks of age i.e. the humane endpoint for untreated KO mice, it remained unclear if treatment with our gene therapy vectors can extend lifespan beyond that age. In our experiments, TG7 proved more effective, achieving higher transgene expression levels and preserving OXPHOS function in the hearts of KO mice. Therefore, we evaluated the long-term efficacy of TG7 at high and low doses (LD and HD, respectively). As before, we injected mice intravenously at 3–4 weeks of age and monitored them over time (Figure 4A). Most of the LD-injected mice succumbed by 18 weeks of age (Figure 4B). Therefore, to avoid losing the final two pairs without being able to carry out any analyses, we euthanized them at 19 weeks of age (Figure 4B). In contrast, only two HD-injected KO mice died prematurely at 22 and 23 weeks (Figure 4B). The remaining HD-injected KO mice showed no overt signs of disease or distress for up to 44 weeks of age, at which point they were sacrificed. Thus, high-dose TG7 injection quadrupled the lifespan of *Lrpprc* KO mice, demonstrating the long-term benefit of this treatment.

**Figure 4.**
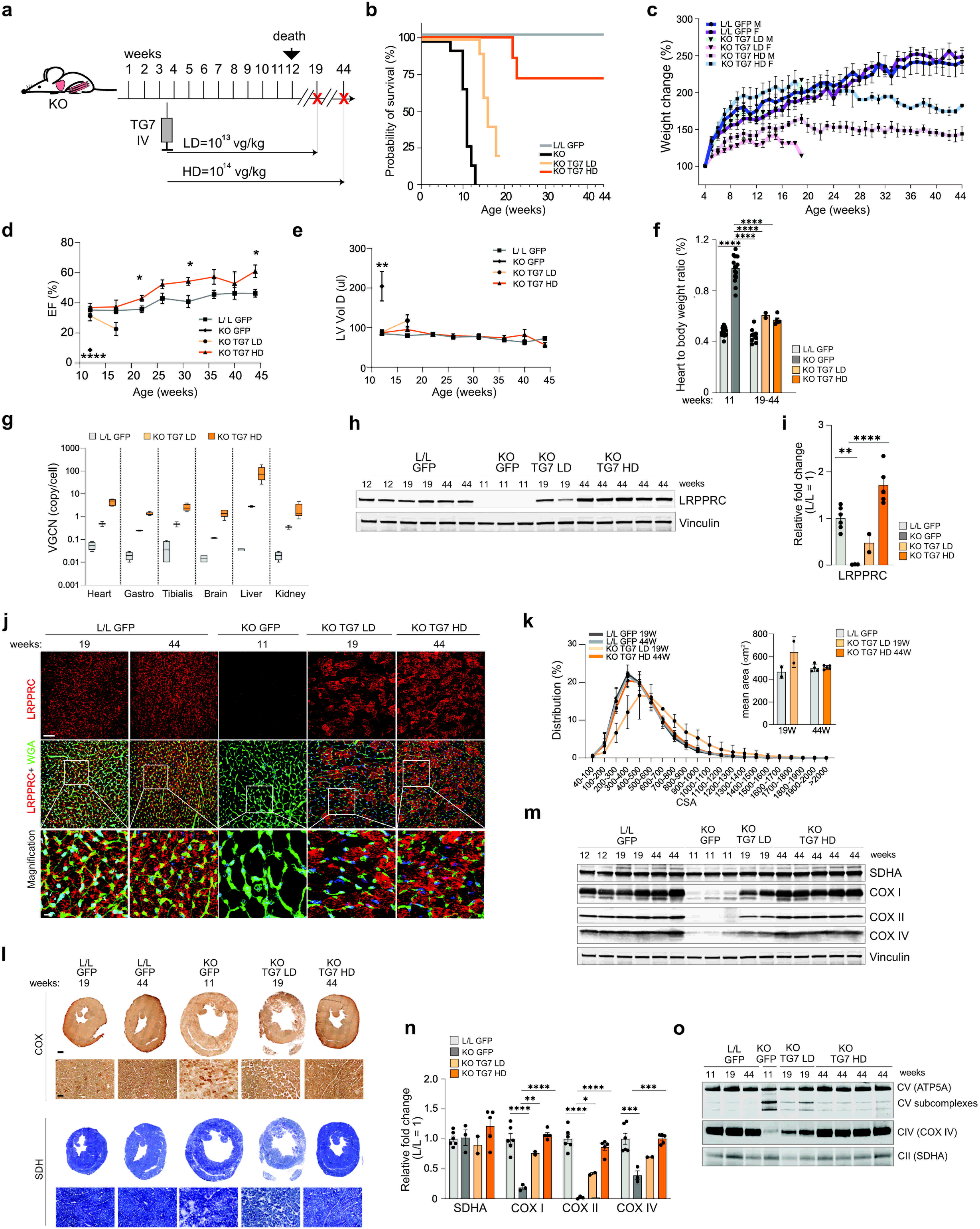
Long-term hLRPPRC expression preserves cardiac structure and function and prevents premature death in KO mice. A) Experimental design for evaluating the long-term therapeutic benefit of TG7 treatment. At 3–4 weeks of age, *Lrpprc* KO mice were injected intravenously with rAAV9-TG7 at either a low dose (LD, 10¹³ vg/kg) or a high dose (HD, 10¹⁴ vg/kg). L/L received TG7-GFP at the corresponding doses. Red crosses indicate euthanasia time points. B) Kaplan–Meier survival analysis of the indicated experimental groups. L/L GFP, n=10; KO TG7 LD, n = 10; KO TG7 HD, n = 7. Survival of untreated KO mice (Figure S1C) is shown for reference. C) Relative body weight change of mice from the indicated experimental groups. For males/females: L/L GFP, n = 5/5; KO TG7 LD, n = 7/3; KO TG7 HD, n = 4/3. D) Echocardiographic analysis of cardiac ejection fraction (EF) in the indicated experimental groups. Measurements from male and female mice were pooled. L/L GFP, n = 15 at 12 and 17 weeks, n = 12 at 22 weeks, n = 11 at 26 weeks, and n = 10 at 31– 44 weeks of age; untreated KO, n = 2 at 12 weeks of age; KO TG7 LD, n = 10 at 12 weeks and n = 4 at 17 weeks; KO TG7 HD, n = 7 at 12-22 weeks, n = 5 at 26–44 weeks. Welch’s t-test: *,p < 0.05; ****, p<0.0001. E) Echocardiographic analysis of left ventricular diastolic volume (LV VolD) in the indicated experimental groups. Sample sizes were as in D. Welch’s t-test: **, p < 0.01. F) Heart weight-to-body weight ratio of mice from the different experimental groups. GFP-injected L/L and KO mice at 11 weeks (Figure 2C) are shown for reference. L/L GFP, 19 and 44 weeks old, n = 8; KO TG7 LD, 19 weeks of age, n = 2; KO TG7 HD, 44 weeks of age, n = 5. One-way ANOVA test with Dunnett’s correction: ****, p<0.0001. G) VGCN in different tissues from TG7 LD- and TG7 HD-injected mice. L/L GFP, used as a negative control, n = 4; KO TG7 LD, n = 2; KO TG7 HD, n = 4-5. H) Western blot analysis of hLRPPRC protein steady-state levels in heart homogenates from the different groups. Vinculin was used as a loading control. Mouse age is indicated in weeks. I) Quantification of the immunoblots shown in H. L/L GFP n = 6; KO GFP, n = 3; KO TG7 LD, n = 2; KO TG7 HD, n = 5. One-way ANOVA test with Dunnett’s correction: **,p<0.01; ****, p<0.0001. J) Representative confocal images showing cardiomyocyte transduction and hLRPPRC expression in male mice from the indicated treatment groups. WGA was used to visualize cardiomyocyte membranes. Scale bar: 50 µm; magnification, 10 µm. Corresponding analyses of female mice are shown in Figure S4A. K) Cardiomyocyte CSA distribution in mouse hearts from the different treatment groups. The bar plot shows the mean area. L/L GFP, n = 2-3 at 19 weeks, n = 4-6 at 44 weeks; KO TG7 LD, 19 weeks of age, n = 2; KO TG7 HD, n = 5. L) Representative images of COX/SDH histochemistry in heart sections from male mice in the different treatment groups. Mouse age is indicated in weeks. Scale bar: whole section image, 500 µm; magnified area, 50 µm. Corresponding analyses of female mice are shown in Figure S4B. M) Western blot analysis of SDHA, COXI, COXII, and COXIV proteins in heart homogenates from the different experimental groups. Vinculin was used as a loading control. Mouse ages are indicated in weeks. N) Quantification of the immunoblots shown in M. L/L GFP, n = 6; KO GFP, n = 3; KO TG7 LD, n = 2; KO TG7 HD, n = 5. One-way ANOVA test with Dunnett’s correction: *, p<0.05; **,p<0.01;***, p<0.001; ****, p<0.0001. O) BN-PAGE analysis of OXPHOS complexes II, IV and V in heart mitoplast extracts from the mice in the different experimental groups. Mouse ages are indicated in weeks.

In addition to changes in body weight, we also monitored cardiac morphology and function in treated mice throughout the experiment. The body weight of HD-injected KO mice continued to increase up to 18-24 weeks of age at different rates between treated male and female KO mice, with treated males gaining weight more rapidly than treated female mice. After reaching a peak, body weight remained stable for both sexes, but lower than that of L/L control mice (Figure 4C). Cardiac function and morphology were monitored by longitudinal echocardiography starting at 12 weeks of age (Figure 4D-E). At this age, ejection fraction and left ventricular volume of L/L controls and LD- and HD-injected KO mice were similar but significantly different from those of GFP-injected KO mice affected by cardiomyopathy. By 17 weeks of age, the LD-injected mice exhibited signs of cardiomyopathy, with a reduced ejection fraction and an increased left ventricular volume, whereas no major differences were observed between HD-injected KO mice and L/L control mice up to 44 weeks of age, with the exception of higher ejection fraction in treated KO mice (Figure 4D-E). Moreover, heart-to-body weight ratios in all treated KO mice were comparable to L/L controls, whereas this ratio increased nearly threefold in GFP-injected KO mice (Figure 4F). These findings demonstrate that a single intravenous injection of HD TG7 protects cardiac function and morphology up to 44 weeks of age.

Next, we assessed the biodistribution of TG7, which remained consistent with the short-term study. We observed a dose-dependent distribution, with slightly lower values than in the short-term study, likely reflecting dilution due to organ growth (Figure 4G). Long-term transgene expression in cardiac tissue was also dose-dependent: HD injection resulted in higher hLRPPRC protein levels and near-complete cardiomyocyte transduction (Figures 4H-J and Figure S4A). Cardiomyocyte area and ventricular morphology in HD-injected mice were comparable to controls, whereas LD mice showed enlarged cardiomyocytes and dilated ventricles (Figures 4K, L and Figure S4B). Complex IV activity and subunit abundance increased in a dose-dependent manner, reaching control levels only in HD mice (Figure 4L-N and Figure S4B). BN-PAGE confirmed these findings and additionally showed a more pronounced complex V F1-subcomplexes accumulation in GFP- and LD-injected KO mice than in HD KO mice (Figure 4O). Thus, injection of *Lrpprc* KO mice with a high dose TG7 therapeutic vector achieved stable long-term hLRPPRC expression, preserved OXPHOS abundance, and prevented the development of cardiomyopathy and premature death.

### Long-term correction of cardiomyopathy unmasks slowly-progressive skeletal muscle pathology in Lrpprc KO mice

The observation that high-dose TG7-injected mice stopped gaining weight after a certain age and exhibited decreased locomotor activity (Figure 4C and Figure S5A), despite the apparent absence of cardiomyopathy, pointed to the development of another pathological process. Our short-term experiment showed that complex IV activity and abundance in skeletal muscle were not significantly preserved, even in HD-injected mice, suggesting that muscle pathology may continue to develop over the long term. Therefore, we analyzed the effect of long-term hLRPPRC expression in gastrocnemius and tibialis muscles. In LD-injected KO mice, hLRPPRC was hardly detectable by immunofluorescence and immunoblotting (Figure 5A-C and Figure S5B). In contrast, hLRPPRC expression was detectable in gastrocnemius and tibialis anterior of HD-injected KO mice. However, this was insufficient to preserve complex IV activity or the abundance of COXII and COXIV subunits in either LD- or HD-injected KO mice (Figure 5D-F and Figure S5C). These findings suggest a pronounced complex IV deficiency in skeletal muscle, regardless of injected dose. Next, we proceeded to assess whether prolonged OXPHOS deficiency, in long-surviving treated KO mice, caused tissue damage and ultimately myopathy. Immunofluorescence analysis revealed the presence of occasional central nuclei, hallmarks of myofiber degeneration and regeneration in myopathy, only in HD-injected mice (Figure 5G and Figure S5D). Moreover, fibers with smaller cross-sectional area (CSA) were overrepresented in gastrocnemius and tibialis anterior muscles from these mice. In contrast, short-lived LD-injected mice, presented only a minor shift toward fibers with smaller CSA, and only in tibialis anterior muscle (Figure 5H-I).

**Figure 5.**
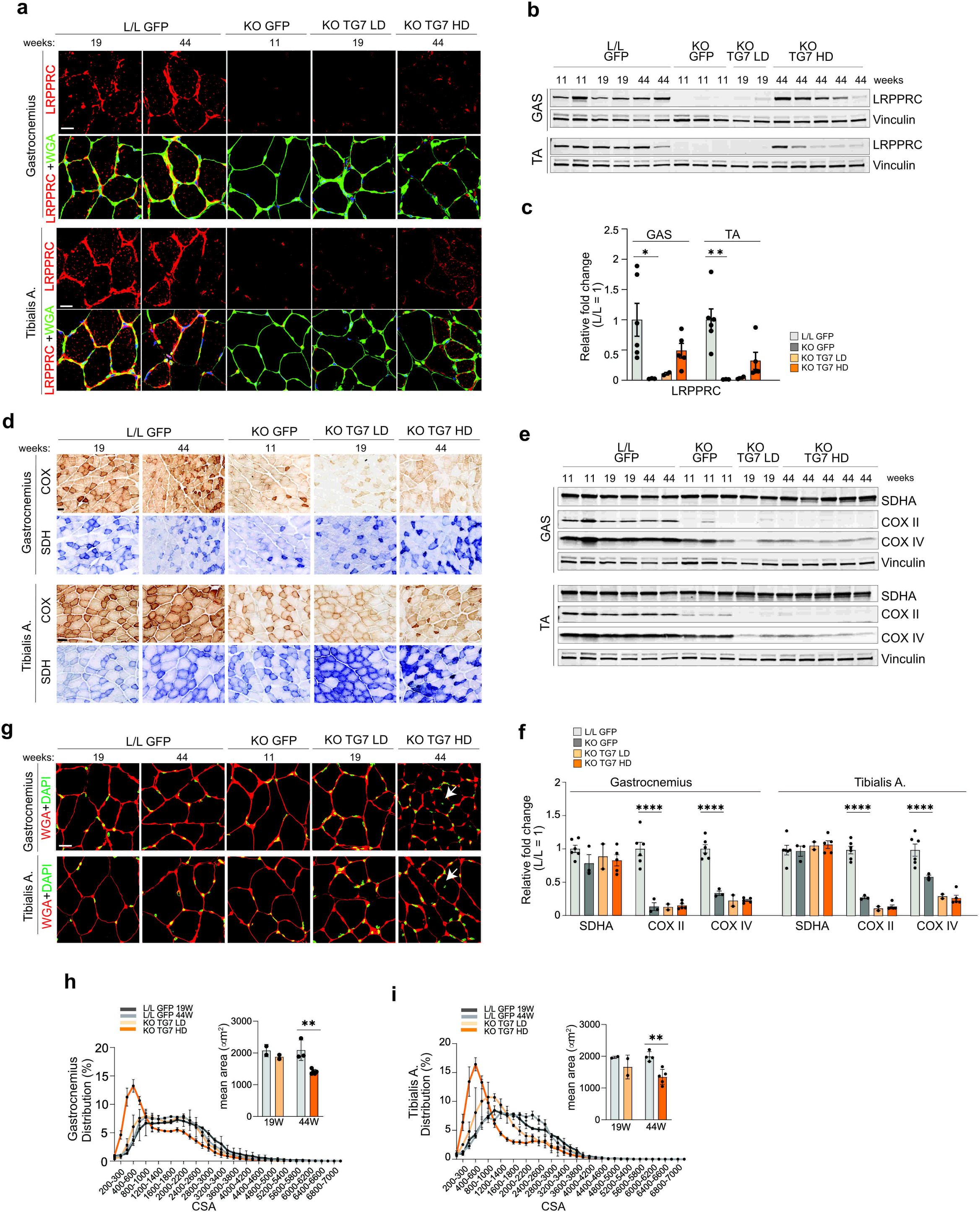
Long-term correction of cardiomyopathy unmasks slowly progressive skeletal muscle pathology in KO mice. A) Representative confocal images showing hLRPPRC expression in gastrocnemius and tibialis anterior muscle sections from male mice from the different groups. WGA was used to visualize muscle fiber membranes. Scale bar: 20 µm. Corresponding analyses of female mice are shown in Figure S5B. B) Western blot analysis of the hLRPPRC protein steady-state levels in gastrocnemius (G, top) and tibialis anterior (TA, bottom) homogenates from the indicated mice. Vinculin was used as a loading control. Mouse age is indicated in weeks. C) Quantification of the immunoblot shown in B. For both muscle types: L/L GFP, n = 6; KO GFP, n = 3; KO TG7 LD, n = 2; KO TG7 HD, n = 5. One-way ANOVA test with Dunnett’s correction: *, p<0.05; **,p<0.01. D) Representative images of COX/SDH histochemical staining in gastrocnemius and tibialis anterior muscle sections of male mice from the indicated experimental groups. Scale bar: 50 µm. Corresponding analyses of female mice are shown in Figure S5C. E) Western blot analysis of SDHA, COXII and COXIV protein steady-state levels in gastrocnemius and tibialis anterior homogenates across the different experimental groups. Vinculin was used as a loading control. Mouse ages are indicated in weeks. F) Quantification of the immunoblots in E. For both muscle types: L/L GFP, n = 6; KO GFP, n = 3; KO TG7 LD, n = 2; KO TG7 HD, n = 5. One-way ANOVA test with Dunnett’s correction: ****, p<0.0001. G) Representative confocal images showing the presence of central nuclei in gastrocnemius and tibialis anterior muscles from male mice in the indicated experimental groups. Scale bar: 20 µm. Corresponding analyses of female mice are shown in Figure S5D. H) Muscle fiber CSA distribution in gastrocnemius muscle. The bar plot shows the mean fiber area. L/L GFP, 19 weeks of age, n = 2, 44 weeks of age, n = 3; KO TG7 LD, 19 weeks of age, n = 2; KO TG7 HD, 44 weeks of age, n = 5. Welch’s t-test: **, p<0.01. I) Muscle fiber CSA distribution in tibialis anterior muscle. The bar plot shows the mean fiber area. L/L GFP, 19 weeks of age, n = 2, 44 weeks of age, n = 4; KO TG7 LD, 19 weeks of age, n = 2; KO TG7 HD, 44 weeks of age, n = 5. Welch’s t-test **, p<0.01.

Together, these results suggest that although KO mice exhibit OXPHOS deficiency without detectable skeletal-muscle pathology at 11 weeks, HD TG7 extends survival by preventing cardiomyopathy but leaves skeletal muscle exposed to prolonged OXPHOS deficiency. These findings highlight tissue-specific thresholds for OXPHOS dysfunction, typical of mitochondrial diseases.

### Sustained mLRPPRC expression has a greater therapeutic benefit in KO mice compared to hLRPPRC

Because of potential differences in RNA-binding specificity between human and murine LRPPRC, and the consequences for the therapeutic potential of our strategy, we tested the efficacy of mouse LRPPRC (mLRPPRC) in preventing disease and extending the survival of KO mice. To this end, we replaced the hLRPPRC sequence in the TG7 plasmid with mLRPPRC, generating TG10 (Figure 6A). Transfection in HeLa cells confirmed that mLRPPRC was expressed and correctly targeted to mitochondria (Figure S6A). As previously described for TG7, rAAV9 viral particles produced from TG10 were administered as a single caudal vein injection in KO mice at 3-4 weeks of age (Figure 6B), and the mice were monitored over time. Among the KO mice injected with LD TG10 some variably succumbed between 13 to 21 weeks of age (Figure 6C). To enable tissue analyses, we sacrificed the last three surviving KO mice at 34 weeks of age (Figure 6C). In contrast, HD TG10-injected mice showed no signs of distress throughout the experiment, except for one mouse that died at 39 weeks of age. The remaining injected mice were sacrificed at 56 weeks of age. However, like HD TG7-injected mice, HD TG10-treated KO mice stopped gaining weight after approximately 18-20 weeks of age (Figure 6D) and exhibited a mild reduction in locomotor activity compared with age-matched control mice (Figure S6B). The heart-to-body-weight ratio of HD and LD TG10-treated KO mice was closer to that of control mice and lower than that of GFP-injected KO mice (Figure 6E). Vector biodistribution analyses showed a pattern similar to TG7, with robust, dose-dependent cardiac transduction (Figure 6F). Immunoblotting confirmed dose-dependent mLRPPRC expression in the heart, although signal intensity appeared lower than that of the endogenous protein (Figure 6G-H) and lower than the hLRPPRC expressed by TG7 (Figure 4H), reflecting the antibody’s higher affinity for the human protein. Immunofluorescence analysis further revealed dose-dependent transduction of cardiomyocytes, with markedly higher transduction efficiency under the HD condition (Figure 6I and Figure S6C). CSA distribution in HD-treated mice was indistinguishable from that of control mice, whereas CSA was increased in LD TG10-injected KO mice, consistent with the presence of cardiomyopathy (Figure 6J). The presence of cardiomyopathy in LD-injected mice was further supported by transverse cardiac sections showing an increased LV area, although this was less pronounced than in GFP-treated KO mice (Figure 6K and Figure S6D). In contrast, LV area in HD-injected mice was similar to controls. Biochemically, mLRPPRC expression preserved complex IV activity and prevented accumulation of complex V subcomplexes already at lower doses (Figure 6K-N and Figure S6D). Together, these data indicate that TG10 expressing the mouse LRPPRC protein outperformed TG7 in prolonging lifespan and preserving OXPHOS function already at LD.

**Figure 6.**
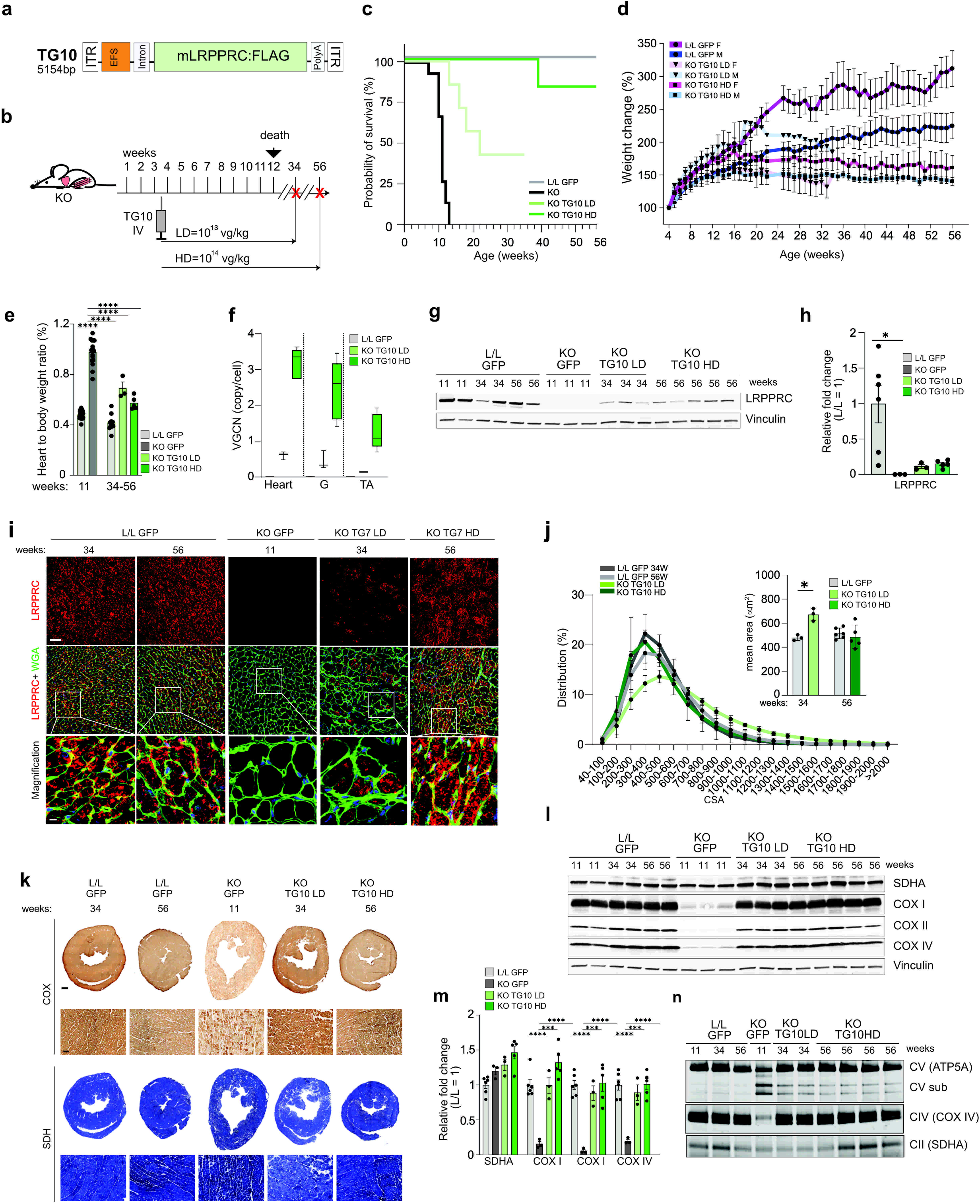
Sustained mLRPPRC expression outperforms hLRPPRC in prolonging survival and preventing cardiomyopathy in KO mice. A) Schematic representation of the TG10 construct expressing mLRPPRC:FLAG. The size between the ITRs is indicated. B) Experimental design for the long-term evaluation of TG10 efficacy. 3–4-week-old *Lrpprc* KO mice were intravenously injected with low-dose (LD, 10¹³ vg/kg) or high-dose (HD, 10¹⁴ vg/kg) rAAV9-TG10. Control mice (L/L) were injected with TG7-GFP at the corresponding LD or HD. Red crosses indicate euthanasia time points. C) Kaplan–Meier survival analysis across the experimental groups. L/L GFP, n = 6; KO TG10 LD, n = 7; KO TG10 HD, n = 6. Survival of untreated KO mice (Figure S1C) is shown for reference. D) Relative body-weight change with age in the different experimental mouse groups. For males/females: L/L GFP, n = 3/3; KO TG10 LD, n = 4/3; KO TG10 HD, n = 3/3. E) Heart weight-to-body weight ratios of mice from the different experimental groups. Data for male and female mice were pooled. L/L GFP, 34 and 56 weeks old, n = 10; KO TG10 LD, 34 weeks old, n = 3; KO TG10 HD, 56 weeks old, n = 5. 11-week-old GFP-injected L/L and KO mice (Figure 2C) are shown for reference. One-way ANOVA test with Dunnett’s correction: ****, p<0.0001. F) VGCN in heart and skeletal muscle of TG10-injected mice. GFP-injected L/L mice were used as a negative control. L/L GFP n = 6; KO TG10 LD, n = 3; KO TG10 HD, n = 5. G) Western blot analysis of mLRPPRC protein steady-state levels in heart homogenates from mice in the different experimental groups. Vinculin was used as a loading control. Mouse age is indicated in weeks. H) Quantification of the immunoblot shown in G. L/L GFP n = 6; KO GFP, n = 3; KO TG10 LD, n = 3; KO TG10 HD, n = 5. One-way ANOVA with Dunnett’s correction: *, p<0.05. I) Representative confocal images showing cardiomyocyte transduction and mLRPPRC expression in males from the indicated experimental groups. WGA was used to visualize cardiomyocyte membranes. Scale bar: 50 µm; magnification 10 µm. Corresponding analyses of female mice are shown in Figure S6C. J) Distribution of cardiomyocyte CSA in the different experimental groups. The mean area is also indicated. L/L GFP, n = 3 at 34 weeks of age, n = 6 at 56 weeks of age; KO TG7 LD, n = 3 at 34 weeks of age; KO TG7 HD, n = 5 at 56 weeks of age. Welch’s t-test: *, p<0.05. K) Representative images of COX/SDH histochemistry in heart transverse sections from male mice in the indicated experimental groups. Mouse age is indicated in weeks. Scale bar: whole section, 500 µm; magnification, 50 µm. Corresponding analyses of female mice are shown in Figure S6D. L) Western blot analysis of SDHA, COXI, COXII and COXIV protein steady-state levels in heart homogenates from the different experimental groups. Vinculin was used as a loading control. Mouse ages are indicated in weeks. M) Quantification of the immunoblot in L. L/L GFP, n = 6; KO GFP, n = 3; KO TG10 LD, n = 3; KO TG10 HD, n = 5. One-way ANOVA test with Dunnett’s correction: ***, p<0.001; ****, p<0.0001. N) BN-PAGE analysis of OXPHOS complexes II, IV and V in heart homogenates from the indicated mouse groups. Mouse ages are indicated in weeks.

Next, we analyzed the expression of the mLRPPRC transgene in skeletal muscle. The mLRPPRC protein was barely detectable by immunofluorescence and immunoblotting in both gastrocnemius and tibialis muscles from LD- and HD-TG10-treated KO mice (Figure 7A-C and Figure S7A). However, complex IV activity and subunits abundance were preserved to a greater extent compared to HD TG7-injected KO mice (Figure 7D-F and Figure S7B). Despite this, occasional central nuclei were present in both muscle types from LD- and HD-treated KO mice (Figure 7G and Figure S7C). Muscle fibers with smaller CSA were more abundant in both treatment groups compared to control mice, suggesting the presence of muscle pathology (Figure 7H-I).

**Figure 7.**
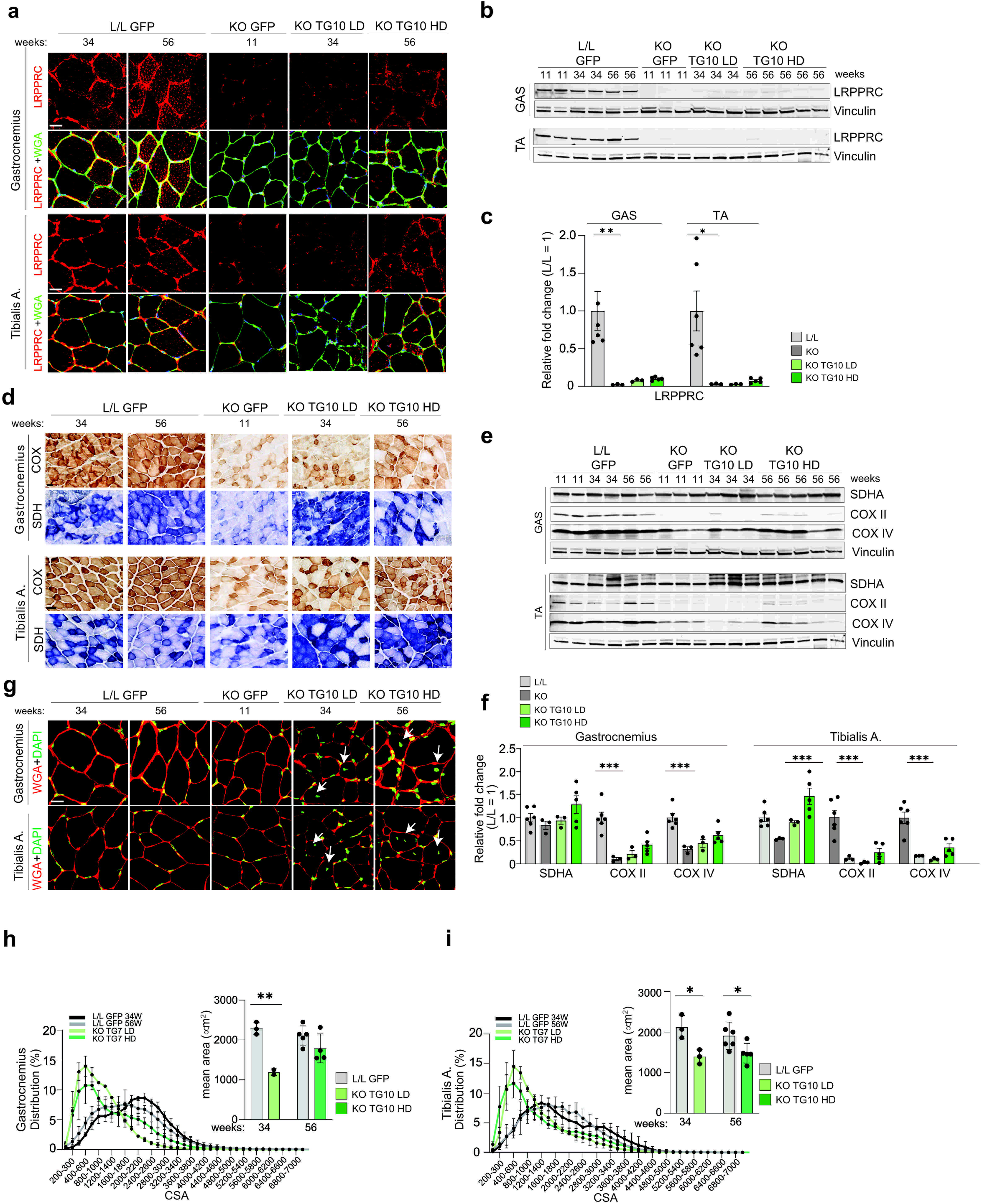
Long-term mLRPPRC expression in skeletal muscle partially preserves OXPHOS function without preventing the development of muscle pathology. A) Representative confocal images showing mLRPPRC expression in gastrocnemius and tibialis anterior muscle sections from males in the different experimental groups. WGA was used to visualize muscle fibers. Scale bar: 20µm. Corresponding analyses of female mice are shown in Figure S7A. B) Western blot analysis of mLRPPRC protein steady-state levels in gastrocnemius (G) and tibialis anterior (TA) homogenates in the different experimental groups. Vinculin was used as a loading control. Mouse age is indicated in weeks. C) Quantification of the immunoblot shown in B. For both muscle types: L/L GFP, n = 6; KO GFP, n = 3; KO TG10 LD, n = 3; KO TG10 HD, n = 5. One-way ANOVA test with Dunnett’s correction: *, p<0.05; **, p<0.01. D) Representative images of COX/SDH histochemistry in gastrocnemius and tibialis anterior muscle sections from the indicated male mouse groups. Scale bar: 50 µm. Corresponding analyses of female mice are shown in Figure S7B. E) Western blot analysis of SDHA, COXII and COXIV protein steady-state levels in gastrocnemius (G) and tibialis anterior (TA) homogenates in the different experimental groups. Vinculin was used as a loading control. Mouse ages are indicated in weeks. F) Quantification of the immunoblot shown in E. For both muscle types: L/L GFP, n = 6; KO GFP, n = 3; KO TG10 LD, n = 3; KO TG10 HD, n = 5. One-way ANOVA test with Dunnett’s correction: ***, p<0.001. G) Representative confocal images showing the presence of central nuclei in gastrocnemius and tibialis anterior muscle from males in the different experimental groups. Scale bar: 20 µm. Corresponding analyses of female mice are shown in Figure S7C. H) Distribution of muscle fiber CSA in gastrocnemius from the indicated experimental groups. The bar plot shows the mean muscle fiber area. L/L GFP, n = 3 at 34 weeks of age, n = 5-6 at 56 weeks of age; KO TG7 LD, n = 2 at 34 weeks of age; KO TG7 HD, n = 4-5 at 56 weeks of age. Welch’s t-test: **, p<0.01 I) Distribution of muscle fiber CSA in tibialis anterior from the indicated experimental groups. The bar plot shows the mean muscle fiber area. L/L GFP, n = 3 at 34 weeks of age, n = 6 at 56 weeks of age; KO TG7 LD, n = 3 at 34 weeks of age; KO TG7 HD, n = 5 at 56 weeks of age. Welch’s t-test: *, p<0.05.

Taken together, these data indicate that mLRPPRC has better therapeutic benefit than hLRPPRC, especially at lower doses. However, it was unable to prevent the development of muscle pathology in treated KO mice.

## Material and Methods

### Vector construction and AAV production

The plasmid pAAV.CMV.PI.EGFP.WPRE.bGH, a gift from Prof. James M. Wilson (Addgene plasmid # 105530 ; http://n2t.net/addgene:105530; RRID:Addgene_105530), was used as the backbone for all the plasmid constructs.

For the trans-splicing dual vector strategy, two plasmids, named L1 and L3, were generated. L1 was constructed by replacing the NotI–XhoI fragment of the pAAV.CMV.PI.EGFP.WPRE.bGH backbone with a sequence encoding the first 20 exons of hLRPPRC, fused at its 3′ end to a splice donor sequence. This cloning step removed the GFP coding sequence together with the WPRE and polyadenylation signal. L3 was generated by replacing the promoter, intron, and GFP cassette of the same backbone with a fragment (AflII–BamHI) containing the remaining 3′ portion of hLRPPRC (last 18 exons), fused at its 5′ end to a splice acceptor sequence and at its 3′ end to a FLAG tag followed by a stop codon.

For the single-vector strategy, several plasmids were generated using extremely compact regulatory elements. Specifically, two ubiquitous promoters were tested: miniCMV (180bp; [45]) and EFS (245bp; [44]) in combination with the same chimeric intron resulting from the fusion of human beta globulin immunoglobulin heavy chain introns (133bp), used in L1 plasmid and the SV40 poly A signal (131bp). Two synthetic double-stranded DNA fragments, each containing one of the above-mentioned promoters, the chimeric intron, the restriction enzyme sites for XbaI, NotI, and SpeI, and the SV40 polyA signal, were synthesized by GeneCust. These fragments were cloned into the pAAV.CMV.PI.EGFP.WPRE.bGH plasmid as an NheI-XhoI fragment, substituting all the regulatory elements except for the two AAV2 ITRs. Next, hLRPPRC:FLAG, mLrpprc:FLAG, and GFP were cloned into these two modified plasmids as NotI-SpeI fragments to generate the following plasmids: TG5 (miniCMV-hLRPPRC:FLAG), TG7 (EFS-hLRPPRC:FLAG), TG10 (EFS-mLrpprc:FLAG), TG5-GFP (miniCMV-GFP), and TG7-GFP (EFS-GFP). rAAV9 particles were produced using an adenovirus-free transient transfection method of HEK293 cell (ATCC) and were purified by CsCl gradient. Titers of rAAV9 particles were determined by two independent methods: RealTime PCR and SDS-PAGE followed by SYPRO Ruby protein gel staining and densitometry.

### Mouse housing, injections and tissue collection

Mouse handling was performed in accordance with French and European legislation on animal welfare (2010/63/EU), and all mouse procedures were approved by the local institutional ethical committee (CEEA34) and the French Ministry for Higher Education and Research (#47432-2024011618239362 v5; #47766-2024032022445779 v1; #47458-2024020914273449 v3). Mice were housed in rooms with a controlled environment (20– 22°C, 45–64% humidity) under a 12 h light/dark cycle and fed ad libitum with a standard diet.

Lrpprc conditionally targeted mice (*Lrpprc^LoxP/LoxP^*), were provided by Prof. Nils-Goran Larsson (Max-Planck for Biology of Ageing, Cologne, Germany), and were maintained in the C57Bl6/N background as previously described in [4]. Transgenic mice expressing *Cre*-recombinase under the creatin kinase promoter (*ckmm-Cre*) were purchased from JAX (#006475) and were backcrossed 10 times to C57Bl6/N mice (Charles River). Heart and skeletal muscle-specific *Lrpprc* knockout mice (KO) were generated by crossing *Lrpprc^LoxP/LoxP^* mice with *ckmm-Cre* mice as described in [4]. The reporter line *Rosa26^tdTomato^* [46] was used to map Cre-recombinase expression and was provided by Dr. Alessandra Pierani (Imagine Institute, Paris, France).

Intravenous injection via the tail vein in 3-4 week-old mice, was used to deliver the different rAAV vectors at the specified dose.

Mice were euthanized by cervical dislocation, and the organs of interest were rapidly collected. For molecular analyses, portions of each organ were snap-frozen in liquid nitrogen. For histological analyses, transverse heart sections were embedded in OCT (Roth) and frozen in liquid nitrogen, while the muscle was snap-frozen in isopentane (Roth) pre-cooled in liquid nitrogen.

### Open field test to determine locomotor activity

The open field test was performed by the same experimenter throughout to ensure consistency across trials. The open field apparatus (29 × 21 cm, white background) was placed in a sound-attenuated room in the mouse facility. Each mouse was individually placed in the center of the arena, and its spontaneous behavior was recorded for 2 or 5 minutes using a smartphone camera positioned above the setup. After the test, each mouse was returned to its home cage. The arena was thoroughly cleaned between animals to remove residual odor cues. Video recordings were subsequently analyzed using Kinovea (version 0.9.5) to track and quantify movement parameters as previously described [47].

### Echocardiography

Echocardiography was performed in anesthetized mice using a Vevo 2100 high-resolution ultrasound system (VisualSonics, Toronto, Canada) with a 40 MHz probe (MS-550). Mice were anesthetized with 3.5% isoflurane (induction), which was then maintained with 1.5% isoflurane. Next, mice were depilated in the thoracic region and placed in supine position on a dedicated heating platform, allowing monitoring of ECG, temperature, and respiratory frequency. Parasternal long axis views and M modes were recorded, as well as a short axis view at the papillary muscle level. Ejection Fraction and Cardiac Output (CO) were calculated from both M-mode and B-mode acquisitions of the Left ventricle in long-axis view using the VevoLab Software (Visualsonics).

### DNA extraction and viral genome copy number analyses

DNA was extracted from pulverized tissue using the DNeasy Blood and Tissue kit (Quiagen). Equal concentration of DNA was used to determine viral genome copy number using ddPCR QX200 (Biorad) using specific primers for each target: actin (forward primer: CATCACTATTGGCAACGAGC, reverse primer: CAGGAGTTTTGTCACCTTTAGAT); hLRPPRC (forward primer: CAAAGGAAGATGGCCCAAGTG, reverse primer: GTAGTCTGTTAGCTTCTCCACA), and mLRPPRC (forward primer: AGATCTGGGACAAACTTCAGC, reverse primer: TCTGGTATGTTACTCGATTTGGTT).

### Protein extraction and immunoblotting

For protein extraction, pulverized tissues were lysed in RIPA buffer supplemented with Complete protease inhibitors cocktail (Roche), and incubated on ice for 30 minutes. Samples were subsequently sonicated for 20 seconds using a needle sonicator and centrifuged at 21,000 × g for 20 minutes at 4°C. The resulting supernatants were collected and quantified using Bradford reagent (Thermo Fisher Scientific). 30-60 µg of proteins was loaded on precast 4-15% TGX gel (BioRad) and electroblotted onto LF PVDF membrane using the Trans-Blot Turbo system (BioRad). Immunodetection was performed using the following primary antibodies: LRPPRC (Proteintech, 21175-1-AP), Vinculin (Abcam, Ab129002 or Sigma, V9264), COXI (Abcam, ab14705), COXII (Proteintech, 55070-1-AP), COXIV (Proteintech, 11242-1-AP), VDAC (Proteintech, 55259-1-AP). Fluorescent secondary antibodies were purchased from Li-COR, and visualization was performed using the Odyssey imaging system (LI-COR Biosciences). Protein quantification was performed using Image Studio (v.5.2) software.

### BN-PAGE analyses

Mitoplasts were purified by differential centrifugation after homogenization of pulverized hearts in 10 mM Tris-HCl, pH 7.5, 320 mM sucrose, supplemented with protease inhibitor cocktail (Roche). Membrane proteins were extracted with n-Dodecyl-β-D-maltoside (DDM) as previously described in [48]. Extracts were fractionated on 4-16% NUPAGE native gel (Thermo Fisher Scientific) and transferred to LF PVDF membrane (Millipore). Immunodetection was performed using the following primary antibodies: SDHA (Abcam, ab14715), COXIV (Proteintech, 11242-1-AP) and ATP5A (Abcam, Ab14748). Fluorescent secondary antibodies were purchased from LI-COR and visualization was performed using the Odyssey imaging system (LI-COR Biosciences).

### Immunofluorescence

14-μm-thick sections from tissues were prepared using a cryostat maintained at −20°C (Leica, CM3050 S). Tissue sections from control and treated mice from the same sex were placed in the same glass slide in order to perform the experiments using the same conditions. Sections were either used immediately or stored at −80°C until analysis. Frozen heart and skeletal muscle cryosections were air dried for 15 min and then fixed with 4% paraformaldehyde in PBS for 20 min in the dark. Next, sections were permeabilized with 0.3% Triton X-100 in PBS for 10 min and blocked either with a blocking solution containing 0.5% bovine serum albumin (BSA) and 2% goat serum or with a Mouse on Mouse IgG blocking reagent (Vector Laboratories) according to the manufacturer’s instructions for 1 h. Following a quick wash with PBS, sections were incubated with primary antibodies against LRPPRC (Proteintech, 21175-1-AP); Tom20 (Santa Cruz, sc-17764) or mCherry (Cliniscience, AB0040-200) diluted in a PBS solution containing 2% goat serum and 0.5% BSA overnight at 4°C. After four washes with PBS, Alexa Fluor-conjugated secondary antibodies (Thermo Fisher Scientific) were applied for 1 hour at room temperature. Alexa Fluor 488-conjugated wheat germ agglutinin (WGA) (Thermo Fisher Scientific) was used to stain plasma membranes. Coverslips were mounted using anti-fade mounting medium (Thermo Fisher Scientific) containing DAPI dye for nuclear staining. Images were acquired with a Leica SP5 confocal microscope using a 40X objective. The brightness and contrast of the acquisitions were adjusted for a better rendering using Fiji software [49].

### Cytochrome c oxidase (COX) and succinate dehydrogenase (SDH) histochemistry

To assess respiratory chain dysfunction in tissue sections, heart and muscle sections were independently stained for cytochrome c oxidase (COX) and succinate dehydrogenase (SDH) activity. Briefly, frozen transverse sections were air-dried for 15 minutes and incubated at 37°C in freshly prepared COX staining solution containing 20 mg/mL catalase (Sigma), 2 mg/mL reduced cytochrome c (Sigma), and 1 mg/mL 3,3′-diaminobenzidine (DAB, Sigma) in 50 mM sodium phosphate buffer, pH 7.4. Sections were then rinsed three times in PBS for 10 minutes each. For SDH staining, sections were incubated in a freshly prepared solution containing 1.5 mg/mL nitroblue tetrazolium (NBT, Sigma), 130 mM sodium succinate, and 0.2 mg/mL phenazine methosulfate (PMS, Sigma) in 0.05 M phosphate buffer, pH 7.6. Sections were then dehydrated through graded alcohols and xylene before mounting. Stained sections were scanned at least 24 hours later using a NanoZoomer 2.0-HT slide scanner (Hamamatsu) and visualized in NDPview2 software.

### Quantification of cardiomyocyte and skeletal muscle fiber cross-sectional area

Cross-sectional area (CSA) was quantified by measuring the area of individual cardiomyocytes or skeletal muscle fibers in whole heart or muscle cross-sections stained with wheat germ agglutinin (WGA). Digital images were acquired using a NanoZoomer 2.0-HT scanner (Hamamatsu). Heart and muscle transverse tissue sections were manually annotated using QuPath [50], and within the annotations, cardiac myocytes and muscle fibers were segmented using the Cellpose2 [51] cyto2 pretrained model. Finally, the area of each segmented fiber was measured.

### Cell culture, transfection, and immunofluorescence

HeLa cells were seeded onto 13-mm coverslips, transfected with Lipofectamine (Thermo Fisher Scientific) and fixed the following day with 4% PFA in PBS for 20 minutes in the dark. Cells were then permeabilized and blocked with a solution containing 0.1% Triton X-100 and 0.5% bovine serum albumin (BSA) in PBS for 1 hour at room temperature. Primary antibodies anti-FLAGM2 (Sigma, F1804), anti-TOM20 (Proteintech, 11802-1-AP) were diluted in the blocking solution and applied overnight at 4°C. After three washes with PBS, Alexa Fluor-conjugated secondary antibodies (Thermo Fisher Scientific) were added for 1 hour at room temperature in the dark. Coverslips were mounted using anti-fade mounting medium (Thermo Fisher Scientific) containing DAPI dye for nuclear staining. Images were acquired using a Leica SP5 confocal microscope equipped with a 63× objective lens. Brightness and contrast adjustments were performed in Fiji.

## Discussion

LRPPRC-linked disease is incurable and fatal. In this study, we provide the first *in vivo* proof-of-concept for AAV-mediated GRT for this disorder. Systemic delivery of compact rAAV9-hLRPPRC vectors in presymptomatic mice preserved cardiac OXPHOS biogenesis, prevented the development of fatal dilated cardiomyopathy and markedly extended survival in heart- and skeletal muscle-specific *Lrpprc* knockout mice. These findings demonstrate that hLRPPRC expression can preserve mitochondrial function in the heart, an LRPPRC-disease-targeted tissue.

A major challenge in developing an AAV-based gene therapy for LRPPRC was the size of its coding sequence, which exceeded the packaging capacity of classical AAV vectors carrying conventional regulatory elements. In our initial trans-splicing dual-AAV strategy, administered at a combined dose of 2 x 10^13^ vg/kg, only a limited number of cardiomyocytes were successfully transduced. Although reconstituted hLRPPRC protein reached immunodetectable levels, this approach preserved complex IV activity and improved survival only modestly. These findings are consistent with the limitations of dual-AAV strategies, which require efficient co-transduction of the same cell and successful reconstitution of a functional expression cassette, reducing overall transduction and expression efficiency compared with single-vector approaches [52]. In contrast to the dual-vector approach, an optimized single-vector design based on compact regulatory elements and elimination of nonessential sequence elements (e.g., WPRE, [53]), produced substantially higher cardiomyocytes transduction and transgene expression which markedly improved efficacy. These experiments suggest that LRPPRC expression must exceed a protein level threshold in a large number of cardiomyocytes to achieve therapeutic benefit. While the minimal level of hLRPPRC expression for functional restoration or maintenance of mtDNA expression has yet to be determined, maximizing transduction efficiency of hLRPPRC expression was critical to achieve therapeutic benefit in our experimental setting.

Another challenge in developing therapies for LRPPRC-linked disease was the lack *of in vivo* models that recapitulate its multisystemic nature while providing clear, quantifiable therapeutic endpoints. The heart- and-skeletal muscle *Lrpprc* KO model used here does not fully reproduce the multisystemic nature of the human disease, particularly its neurological manifestations. Other available mouse models also have constraints: whole-body and brain-specific *Lrpprc* knockout mice are embryonically lethal [4, 54], precluding postnatal therapeutic assessment in these settings; liver-specific *Lrpprc* knockout mice develop tissue abnormalities, but do not show reduced lifespan, precluding survival-based preclinical efficacy studies [16]. In this context, the heart- and skeletal muscle-specific knockout model offers a disease-relevant system with survival-limiting cardiomyopathy and measurable, reproducible functional, histological, and biochemical endpoints. Thus, although this model is insufficient to evaluate correction of neurological and hepatic dysfunction in *LRPPRC*-linked disease, it provides a robust platform to test whether AAV-mediated LRPPRC replacement can prevent fatal OXPHOS-driven cardiomyopathy *in vivo*. Furthermore, using ubiquitous promoters and a broad tissue-tropism capsid, such as AAV9, combined with biodistribution and transgene expression analyses in other tissues, may provide valuable information on the potential of our GRT approach to target multiple organs.

The strongest therapeutic benefit of our GRT was observed in the heart following high-dose systemic AAV administration. In untreated or GFP-treated KO mice, loss of *Lrpprc* resulted in severe complex IV deficiency, accumulation of complex V F1 subcomplexes, cardiomyocyte enlargement, ventricular dilation, impaired cardiac function, and early death. Systemic delivery of HD hLRPPRC prevented progression of cardiac pathology and prolonged survival, whereas LD treatment was considerably less effective. Our findings contrast a previous report using a Friedreich’s ataxia mouse model, in which transduction of approximately half of the cardiomyocytes was sufficient to correct cardiac dysfunction [55]. In our model, systemic administration of LD hLRPPRC transduced a substantial proportion of cardiomyocytes, albeit fewer than the HD treatment. Nevertheless, while sufficient to modestly prolong survival, this level of transduction did not prevent eventual development of fatal cardiomyopathy. Thus, in our setting, the proportion of transduced cardiomyocytes alone does not predict therapeutic efficiency, since, as mentioned above, hLRPPRC expression in individual cardiomyocytes must reach levels sufficient to support mtDNA expression. Therefore, moving forward, it will be important to determine more precisely the relationship between the fraction of transduced cardiomyocytes and hLRPPRC expression levels sufficient to achieve a lasting therapeutic benefit.

*Lrpprc* inactivation in our KO model occurs during embryogenesis. However, it takes at least 4 weeks after birth for LRPPRC protein depletion and complex IV deficiency, while heart function starts to be affected after another 4 weeks, as determined by ECG analyses [4]. Our results show that re-introducing LRPPRC in pre-symptomatic mice can reverse the complex IV defect and prevent development of cardiomyopathy. Going forward, it will be important to determine whether rAAV9-hLRPRPC administration in early-symptomatic mice (e.g. ∼7-8 weeks of age to allow for sufficient time for viral uptake and transgene expression) can similarly reverse the already present biochemical defects, prevent or arrest cardiomyopathy development, and extend lifespan.

Notably, in skeletal muscle, hLRPPRC expression also occurred before detectable tissue damage, but GRT was less effective at preserving complex IV abundance and function. VGCN measurements and rAAV9-mediated protein expression in treated KO mice confirmed skeletal muscle transduction, particularly at the high dose, although to a lesser extent than in the heart. Consequently, hLRPPRC expression was insufficient to fully preserve complex IV activity and subunit abundance. In long-term survivors, persistent OXPHOS deficiency was associated with mild myopathic changes, including smaller muscle fibers and occasional centrally nucleated fibers. These findings suggest that skeletal muscle tolerated LRPPRC deficiency and the resulting OXPHOS dysfunction longer than the heart, but it inevitably also developed signs of impairment. The cardiac rescue achieved in our study prolonged survival sufficiently to reveal skeletal muscle pathology that would otherwise be masked by the early mortality of untreated mice due to cardiomyopathy. The incomplete skeletal muscle rescue observed in our study is consistent with previous similar GRT studies using both cardiac and skeletal muscle specific knockout mice. In Friedreich’s ataxia models, systemic AAV-mediated frataxin expression was sufficient to prevent or reverse cardiomyopathy and improve survival, but long-term rescue was limited by subsequent development of myopathy in treated mice [39]. Similarly, in a mouse model of Barth syndrome, AAV-mediated *TAZ* replacement efficiently prevented or reversed cardiac dysfunction, but provided only modest improvements in skeletal muscle pathology [36]. Together, these studies suggest that the heart is more readily transduced than skeletal muscle following systemic AAV delivery, at least with currently used capsids and ubiquitous promoters. This is corroborated by a GRT study in a skeletal muscle-specific *Ndufs3* knockout model, in which an extremely high dose of rAAV9-Ndufs3 (10^15^ vg/kg) was required to prevent development of myopathy in pre-symptomatic mice or to ameliorate it in post-symptomatic animals [22]. These findings suggest that effective and durable correction of mitochondrial myopathy in skeletal muscle using the classical AAV9 or AAV10 capsid may require higher vector doses than those sufficient for cardiac rescue, likely because correction depends on widespread transduction across large, multinucleated myofibers.

Direct comparison of the human and murine transgenes highlighted important differences. mLrpprc provided a better phenotypic rescue than hLRPPRC, more evident at low doses. The stronger rescue obtained with the murine transgene suggests that murine Lrpprc binds more efficiently to murine mitochondrial mRNAs than human LRPPRC and therefore promotes their translation more effectively. This observation is consistent with our previous report showing that the human LRPPRC–SLIRP complex preferentially binds human RNA targets, whereas the mouse complex preferentially binds mouse targets in vitro [18]. These results suggest that the performance of hLRPPRC in the mouse model may underestimate its potential efficacy in humans.

Differences in the efficiency of rAAV9-mediated hLRPPRC expression in preventing pathology in the heart and skeletal muscle highlight the challenges of developing effective treatments for multisystemic disorders such as MDs. Successful treatment of the life-threatening organ pathology, while lifesaving and beneficial in the near-term, may not be sufficient to achieve long-term clinical benefit, especially if the disease develops also in other organs/systems later in life. Our findings illustrate such a scenario and is analogous to that observed in some mitochondrial disorders where organ transplantation cured a life-threatening pathology but was subsequently followed by the progression of disease in other affected organs [56–58]. Achieving a broad tissue correction may require novel AAV serotypes with wide tissue tropism, ubiquitous promoters, optimized routes of administration and specific time windows for the administration. Because GRT elicits long-lasting immune responses, its re-administration is currently not feasible. Therefore, therapeutic strategies for multisystemic mitochondrial diseases should ideally aim for broad, durable correction across tissues. Future treatments for multisystemic diseases can also employ a multipronged approach incorporating GRT and pharmacological treatment, and, where necessary and feasible, an organ transplant. For now, we have demonstrated the first proof of concept for LRPPRC-linked GRT paving the way for future development in the field.

## Supplementary Figure legends

**Figure S1. The Lrpprc knockout mouse model and evaluation of the trans-splicing dual vector gene replacement strategy**

A) Mapping of ckmm-Cre expression. Top: breeding scheme to generate a reporter model with active tdTomato expression. Bottom: representative confocal images showing reporter expression in striated muscle. tdTomato was detected using α-Cherry antibodies. WGA was used to visualize cardiomyocyte and skeletal muscle fiber membranes. Scale bar: 10 µm.
B) Echocardiographic analysis of ejection fraction (EF) and left ventricular diastolic volume (LV VolD) in L/L (control, n = 4) and KO (n = 3-4) mice at different ages. Welch’s t-test: **,p<0.01; ****, p<0.0001.
C) Kaplan–Meier survival analysis of control mice (L/L, pooled male and female mice, n = 15) and KO mice (pooled male and female, n=15).
D) Schematic representation of the two constructs (L1 and L3) used for the trans-splicing dual vector strategy to express hLRPPRC. Regulatory elements are indicated. SDS: splice donor sequence; SAS: splice acceptor sequence.
E) Experimental design for evaluation of the efficacy of the trans-splicing dual vector strategy. 3–4-week-old KO mice were intravenously injected with L1 and L3 using a total dose of 2×10¹³ vg/kg viral particles. Control mice were injected with GFP-expressing viral particles at the same dose. Red crosses indicate euthanasia time point.
F) Western blot analysis of hLRPPRC, COXII and COXIV protein steady-state levels in mitochondrial extracts from the indicated experimental groups. VDAC was used as a loading control.
G) Representative confocal images showing cardiomyocyte transduction and hLRPPRC expression in the different experimental groups. WGA was used to visualize cardiomyocyte membranes. Scale bar: 50 µm; magnification, 10 µm.
H) Representative images of COX/SDH histochemistry in heart sections from the indicated experimental groups. Scale bar: 50 µm.

**Figure S2. Efficacy of hLRPPRC expression in female KO mice**

A) Western blot analysis of hLRPPRC protein levels in total tissue protein extracts from the heart, gastrocnemius, tibialis anterior, and liver of female mice from the indicated experimental groups. Vinculin was used as a loading control.
B) Representative confocal images of transverse heart sections from 11-week-old female mice demonstrating hLRPPRC expression following administration of the indicated vectors. WGA was used to visualize cardiomyocyte boundaries. Scale bars: main image 50 µm, magnification 10 µm.
C) Distribution of cardiomyocyte cross-sectional area (CSA) in heart sections from female mice in the indicated experimental groups. The bar plot shows the mean muscle fiber area. L/L, n = 12; KO GFP, n = 6; KO TG5 LD, n = 4; KO TG7 LD, n = 4; KO TG5 HD, n = 5, KO TG7 HD, n = 5. One-way ANOVA test with Dunnett’s correction: **, p<0.01; ***, p<0.001; ****, p<0.0001.
D) Representative images of COX/SDH histochemical staining of transverse heart sections for female mice in the different experimental groups. Scale bar: whole section, 500 µm; magnified region, 50 µm.
E) Western blot analysis of the steady-state levels of SDHA, COX I, COX II and COX IV in heart homogenates from females in the indicated experimental groups. Vinculin was used as a loading control.
G) BN-PAGE analysis of OXPHOS complexes II, IV and V in heart mitoplast extracts from female mice in the different experimental groups.

**Figure S3. hLRPPRC expression and OXPHOS function in skeletal muscles**

A) Representative confocal images of skeletal muscle sections showing expression of endogenous mLRPPRC and its colocalization with the mitochondrial marker TOM20. WGA was used to visualize muscle fibers. Scale bar: 20 µm.
B) Western blot analysis of SDHA, COX I, COX II and COX IV protein abundance in tibialis (top) and gastrocnemius (bottom) homogenates from female mice in the different treatment groups.

**Figure S4. Analysis of the effect of long-term hLRPPRC expression female KO mice**

A) Representative confocal images showing cardiomyocyte transduction and hLRPPRC expression in female mice from the indicated treatment groups. WGA was used to visualize cardiomyocyte membranes. Scale bar: 50 µm; magnification, 10 µm.
B) Representative images of COX/SDH histochemistry in heart sections from female mice in the different treatment groups. Mouse age is indicated in weeks. Scale bar: whole section image, 500 µm; magnified area, 50 µm.

**Figure S5. Analysis of the effect of long-term hLRPPRC expression in skeletal muscle of KO mice**

A) Locomotor performance in the open-field test of female and male mice in the different experimental groups. 11-week-old GFP-injected L/L and KO mice (Figure 1E) are shown for reference. L/L GFP, 44 weeks of age, n = 5; KO TG7 HD, 44 weeks of age, n = 5. Mann-Whitney test: **,p<0.01; ****, p<0.0001.
B) Representative confocal images showing hLRPPRC expression in gastrocnemius and tibialis anterior muscle sections from female mice from the different groups. WGA was used to visualize muscle fiber membranes. Scale bar: 20 µm.
C) Representative images of COX/SDH histochemical staining in gastrocnemius and tibialis anterior muscle sections of female mice from the indicated experimental groups. Scale bar: 50 µm.
D) Representative confocal images showing the presence of central nuclei in gastrocnemius and tibialis anterior muscles from female mice in the indicated experimental groups. Scale bar: 20 µm.

**Figure S6. Effect of long-term mLRPPRC expression in hearts from KO mice**

A) Representative confocal images of HeLa cells transiently transfected with the TG10 plasmid, showing mLRPPRC:FLAG expression and its mitochondrial localization. TOM20 was used to detect mitochondria. Scale bar: 10 µm; magnifcation
B) Locomotor performance in the open-field test of female and male mice in the different experimental groups. 11-week-old GFP-injected L/L and KO mice (Figure 1E) are shown for reference. L/L GFP, 56 weeks of age, n = 6; KO TG10 HD, 56 weeks of age, n = 5. Mann-Whitney test: **,p<0.01; ****, p<0.0001.
C) Representative confocal images showing cardiomyocyte transduction and mLRPPRC expression in females from the indicated experimental groups. WGA was used to visualize cardiomyocyte membranes. Scale bar: 50 µm; magnification 10 µm.
D) Representative images of COX/SDH histochemistry in heart transverse sections from female mice in the different experimental groups. Mouse age is indicated in weeks. Scale bar: whole section, 500 µm; magnification, 50 µm.

**Figure S7. Long-term expression of mLRPPRC in skeletal muscle of female KO mice**

A) Representative confocal images showing mLRPPRC expression in gastrocnemius and tibialis anterior muscle sections from female mice in the different experimental groups. WGA was used to visualize muscle fibers. Scale bar: 20µm.
B) Representative images of COX/SDH histochemistry in gastrocnemius and tibialis anterior muscle sections from the indicated female mouse groups. Scale bar: 50 µm.
C) Representative confocal images showing the presence of central nuclei in gastrocnemius and tibialis anterior muscle from female mice in the different experimental groups. Scale bar: 20 µm.

## Acknowledgements

This study was supported by AFM-Telethon # 22251, #24318, DIM (*Domaine d’Intérêt Majeur*) of the Paris Ile-de-France Region grant and The Dillon Fund.

BR and SZ were supported by a fellowship from the AFM-Telethon #23291 and #24199. We are grateful to Nicolas Goudin from the Necker Bioimages Analysis Platform, the Mouse Facilities of Imagine Institute and the Necker campus, and the Histology Core Facility, SFR Necker Inserm US24–CNRS UAR 3633. We also thank ARC for providing access to the NanoZoomer 2.0 HT scanner (Hamamatsu) and Dr. Federico Chiossi for his valuable support with data analysis. Echocardiography was performed at the Life Imaging Facility of Paris Cité University (Plateforme Imageries du Vivant—PIV), supported by France Life Imaging (grant ANR-11-INBS-0006), Région Ile-de-France, SIRIC CARPEM (grant INCa-DGOS-Inserm-ITMO Cancer_18006) and IBISA consortium, and financial support from ITMO Cancer of Aviesan within the framework of the 2021-2030 Cancer Control Strategy, on funds administered by Inserm».

This work is dedicated to the memory of Dr. Orazio Ruzzenente, an inspiring scientist and a loving father, whose enduring support helped sustain this project through its most difficult moments.

