## Supplementary figures and images for "LRPPRC gene replacement therapy preserves cardiac function and prevents premature death in a mouse model of mitochondrial disease"

### FIGS1

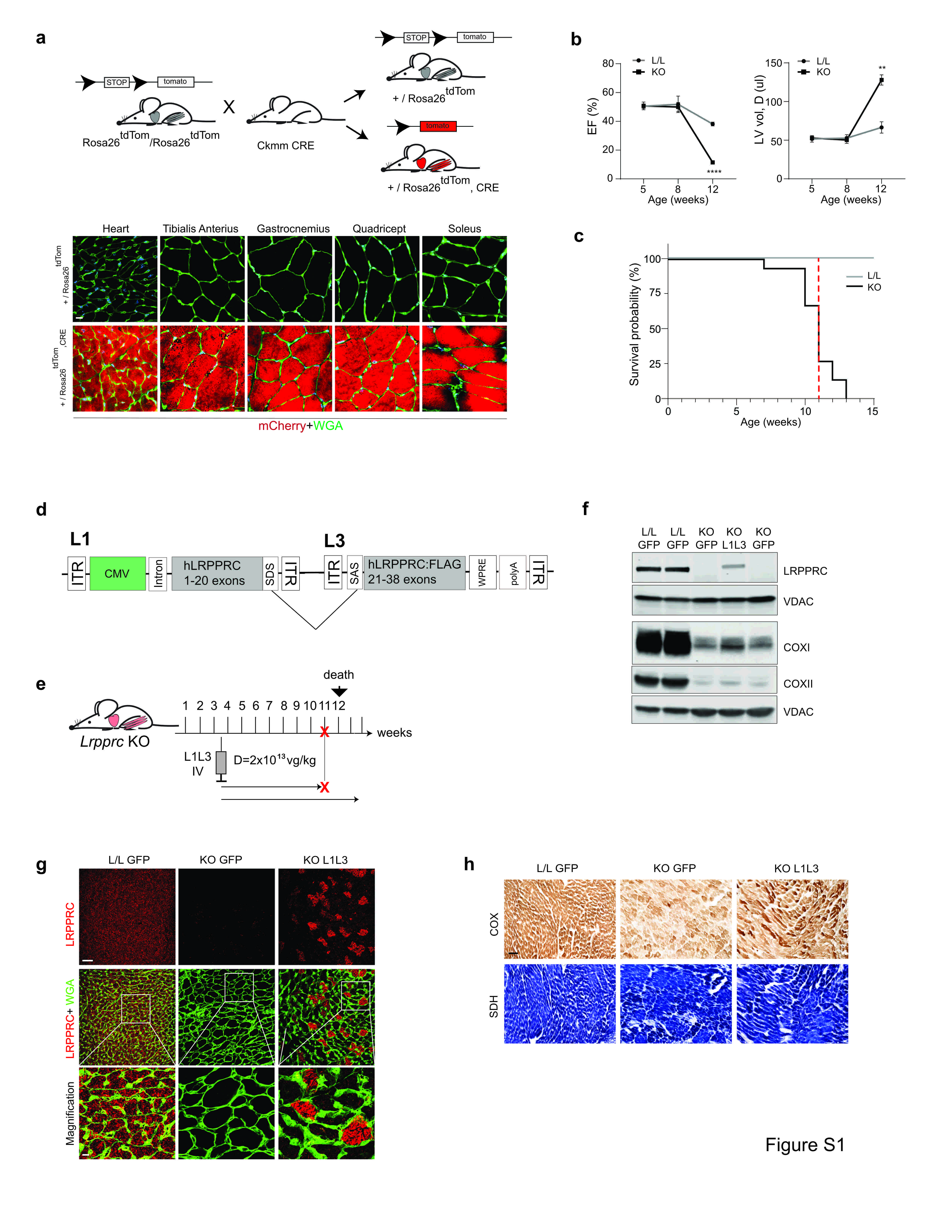

### FIGS2

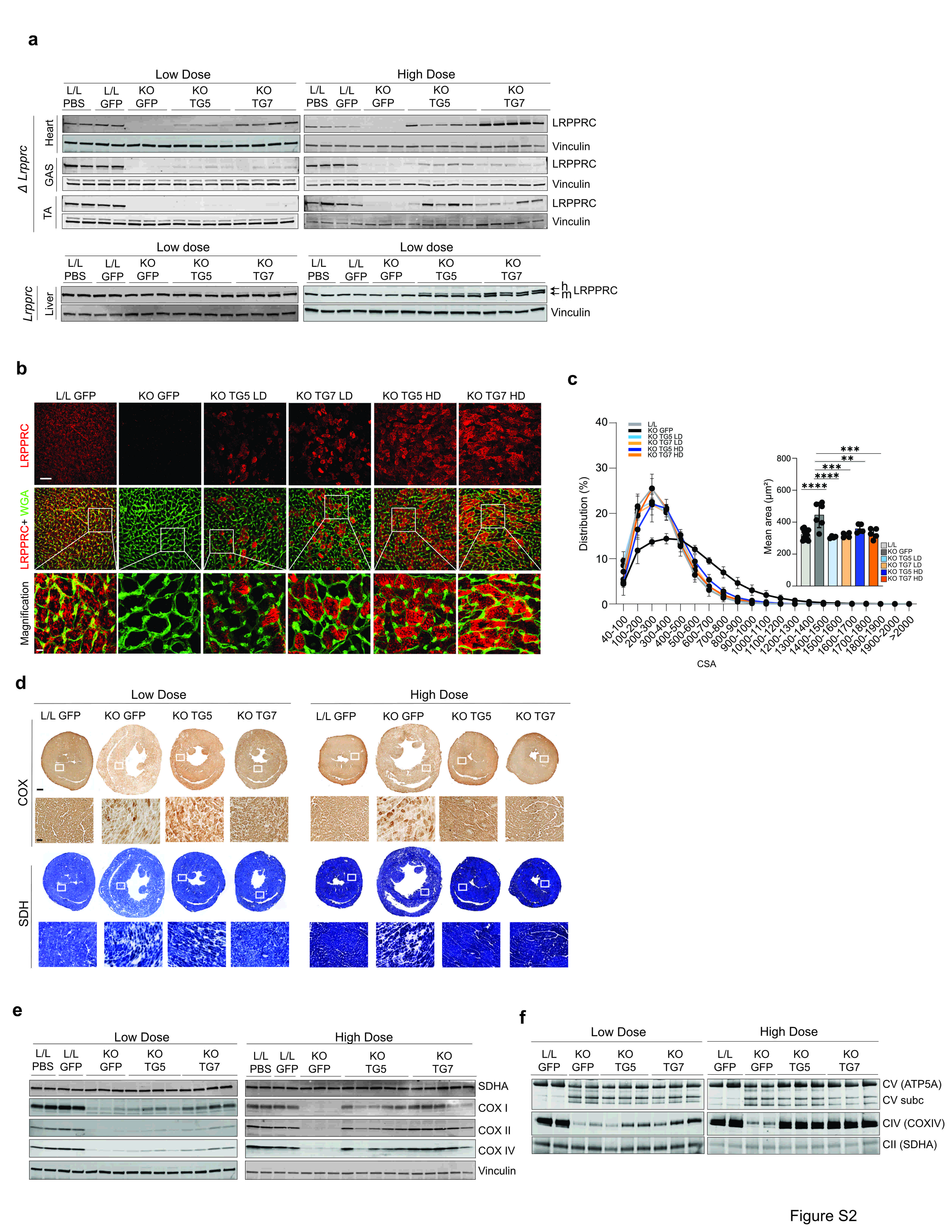

### FIGS3

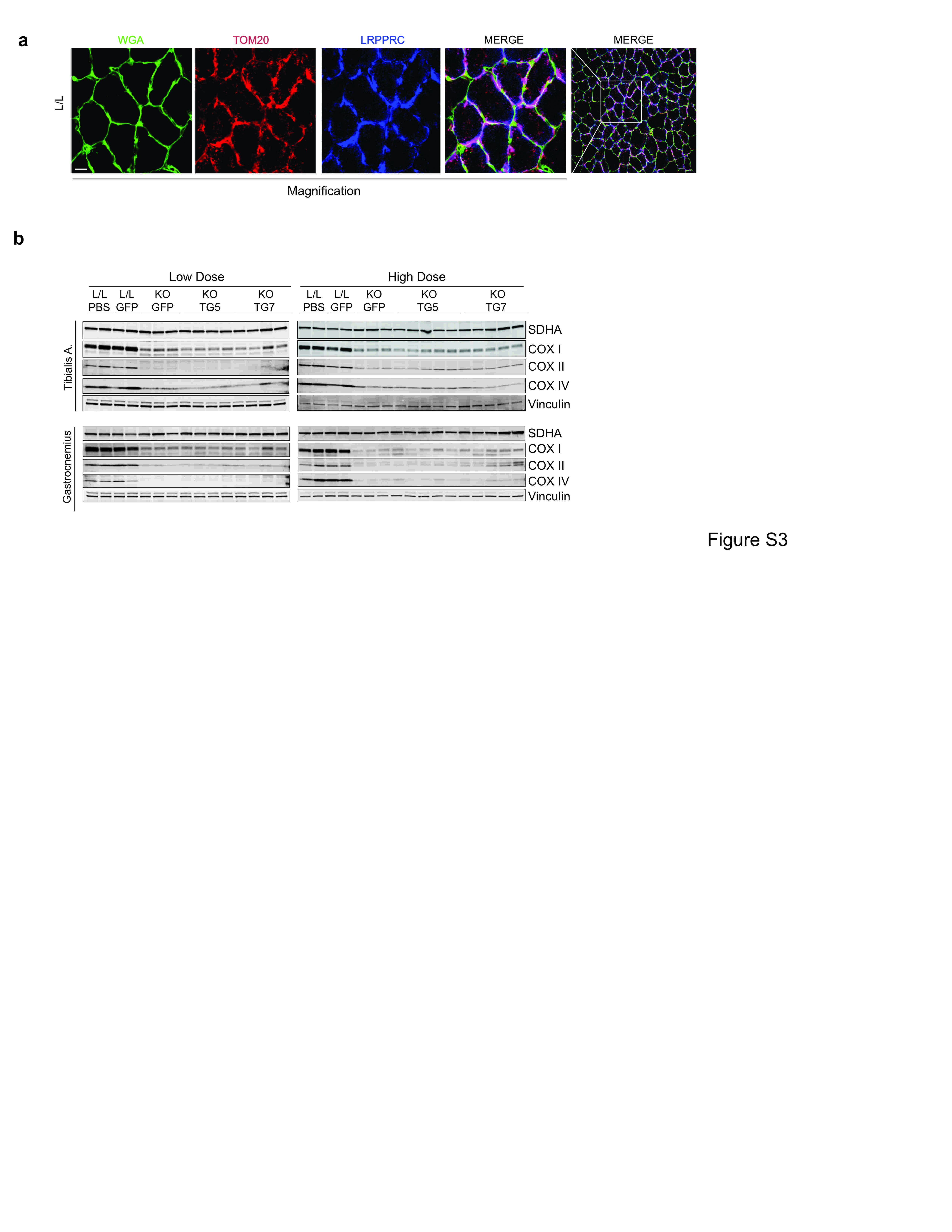

### FIGS4

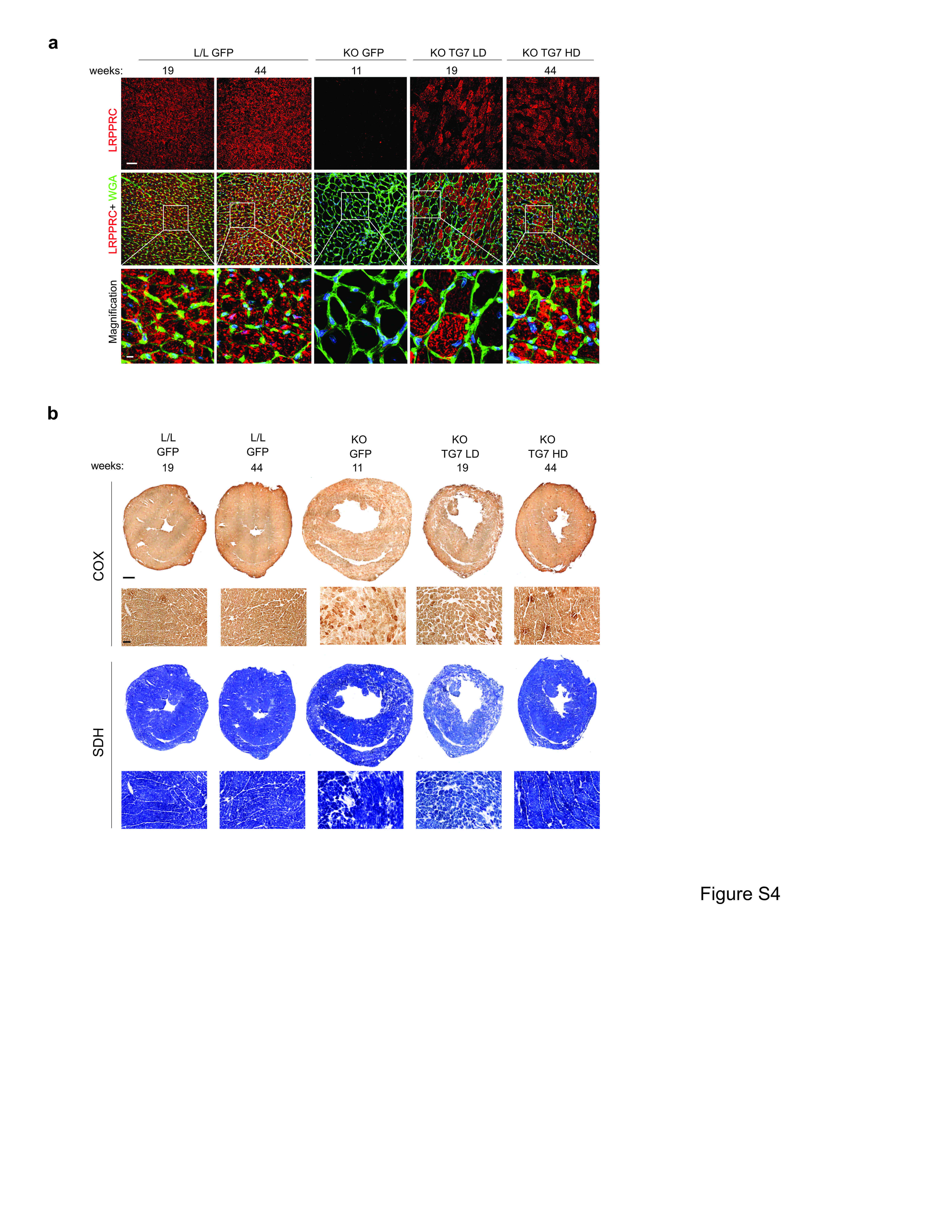

### FIGS5

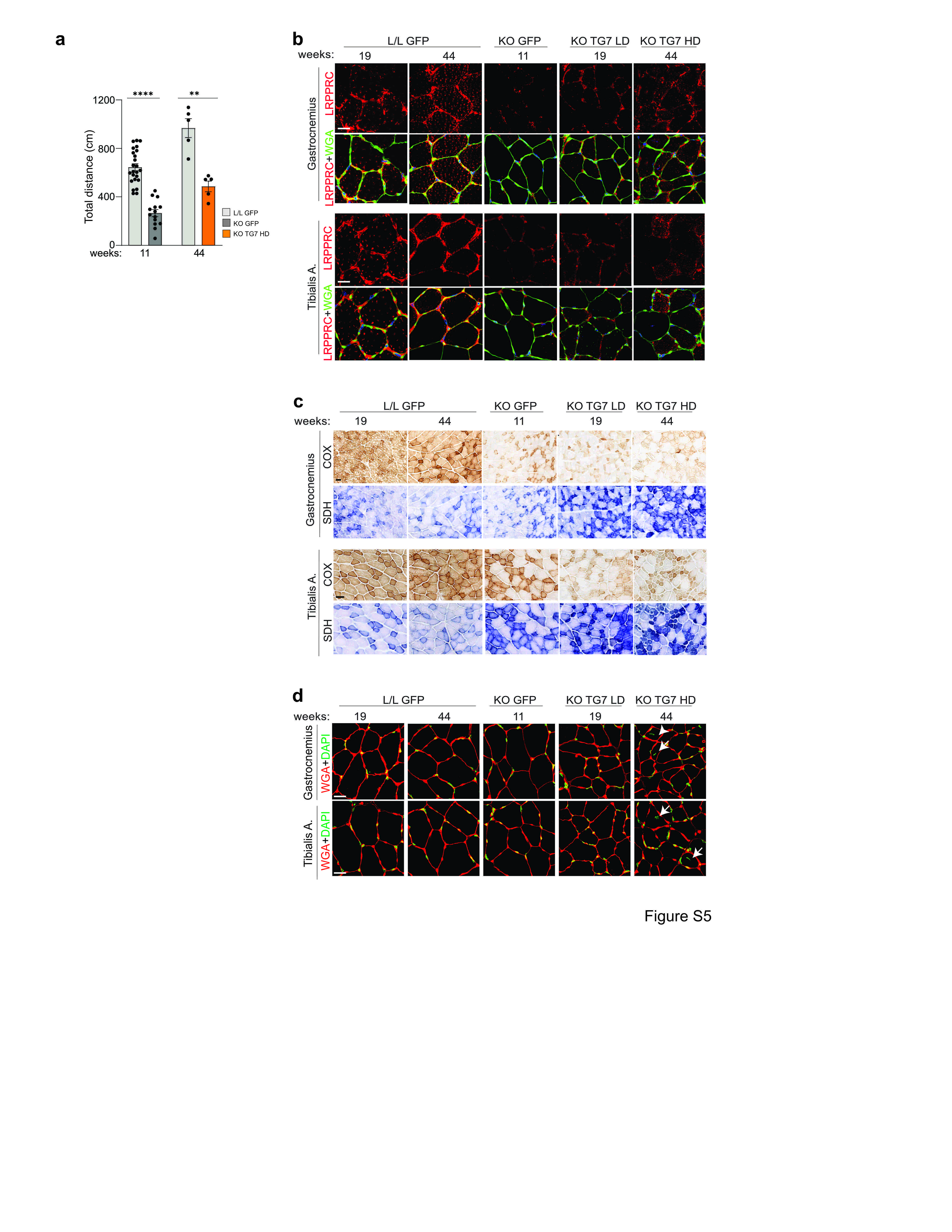

### FIGS6

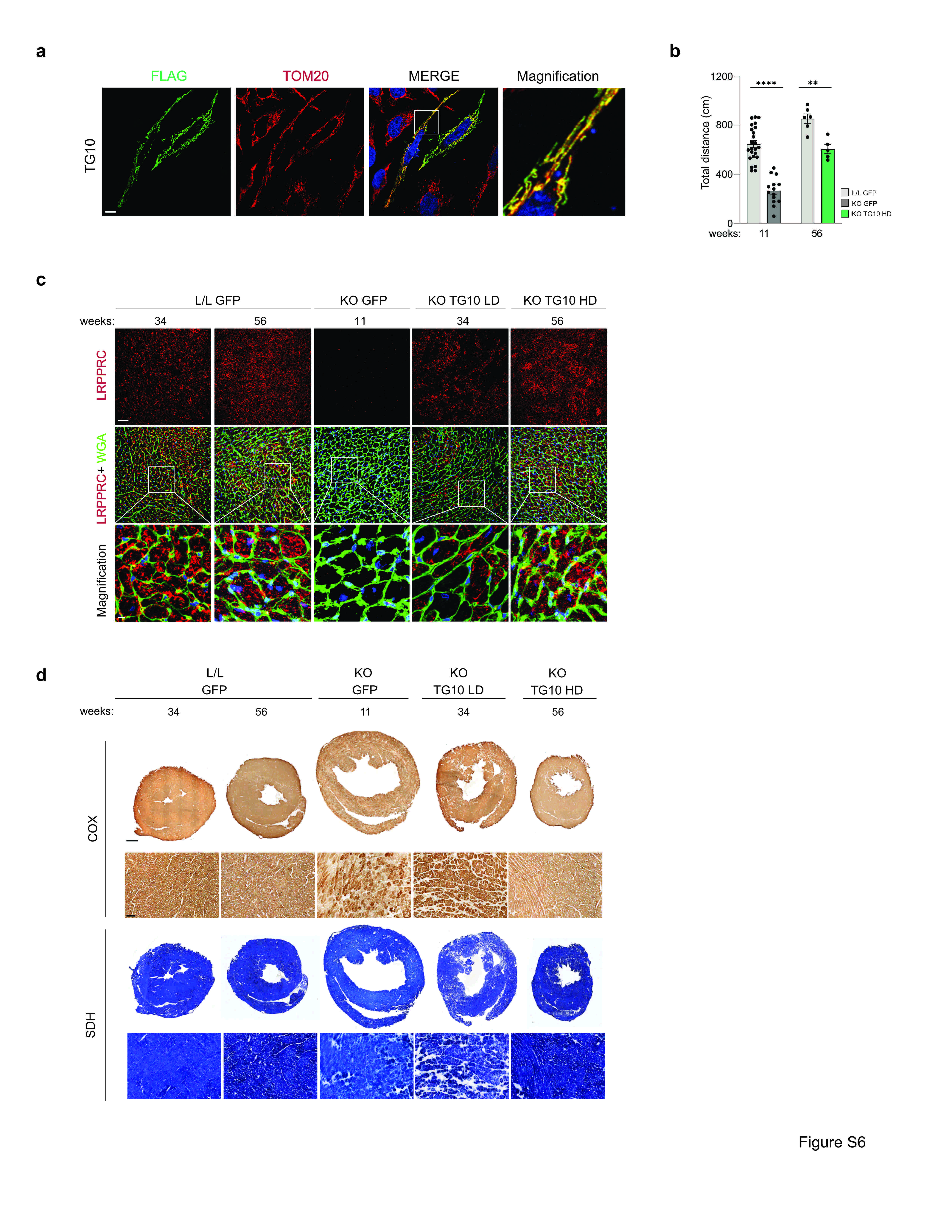

### FIGS7

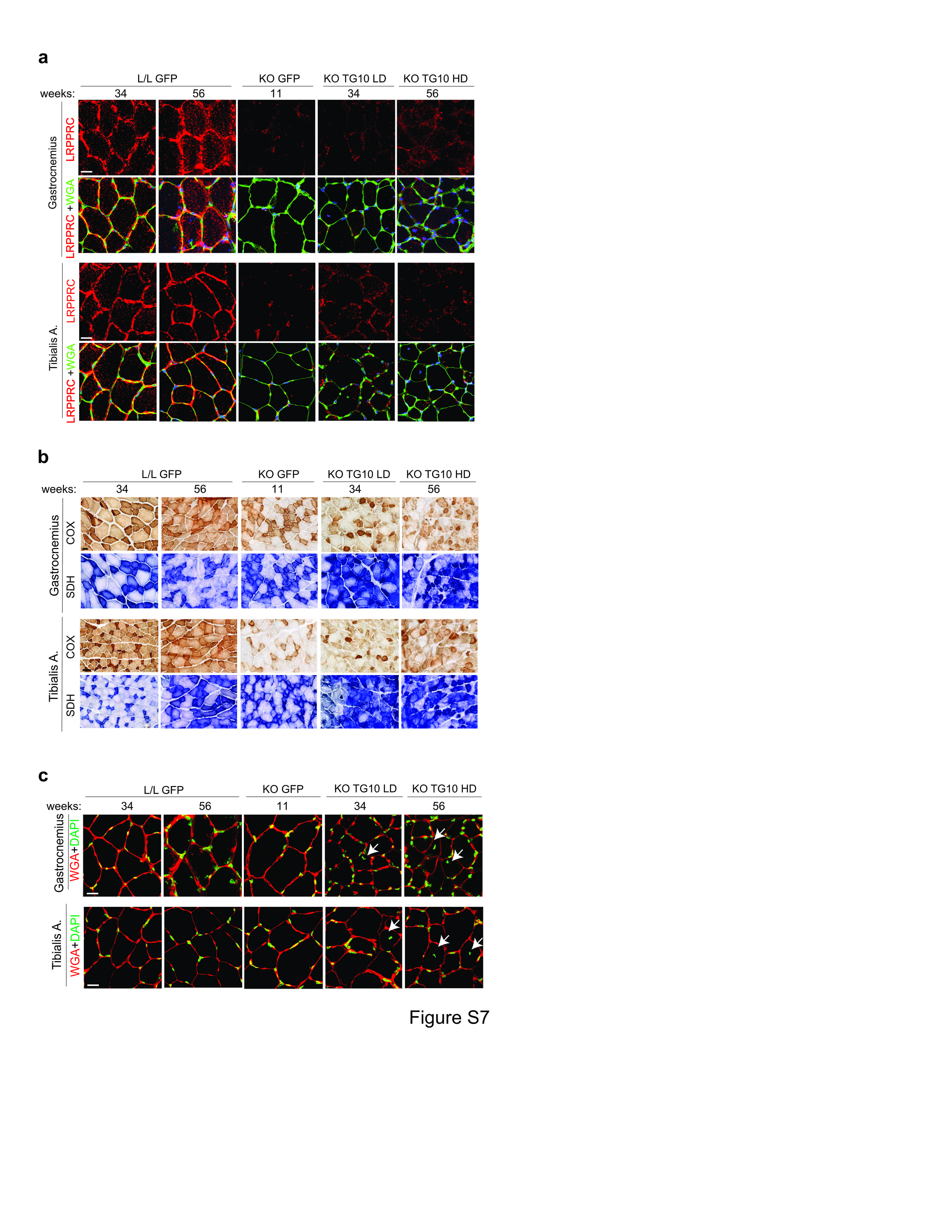
